# Impacts of sequence and structure on pyrrolocytosine fluorescence in RNA

**DOI:** 10.1101/2024.10.12.618031

**Authors:** Taylor L. Coulson, Julia R. Widom

## Abstract

Fluorescence spectroscopy encompasses many useful methods for studying the structures and dynamics of biopolymers. Applications to nucleic acids require the use of extrinsic fluorophores such as fluorescent base analogues (FBAs), which mimic the native bases but have enhanced fluorescence quantum yields. In this work, we use multiple complementary methods to systematically investigate the sequence- and structure-dependence of the fluorescence of the FBA pyrrolocytosine (pC) within RNA. We demonstrate that pC is typically brightest in conformations in which it is base-stacked but not base-paired, properties that distinguish it from more widely used FBAs. This effect is strongly sequence-dependent, with adjacent adenosine and cytidine residues conferring the greatest contrast between stacked and unstacked structures. Structural heterogeneity was resolved in single-stranded RNA and fully complementary and mismatched double-stranded RNA using time-resolved fluorescence measurements and fluorescence-detected circular dichroism spectroscopy. Double-stranded contexts are distinguished from single-stranded contexts by the presence of inter-strand energy transfer from opposing bases, while base-paired pC is distinguished by its short excited state lifetime. This work will enhance the value of pC as a structural probe for biologically and medicinally significant RNAs by guiding the selection of labeling sites and interpretation of the resulting data.

**GRAPHICAL ABSTRACT.**
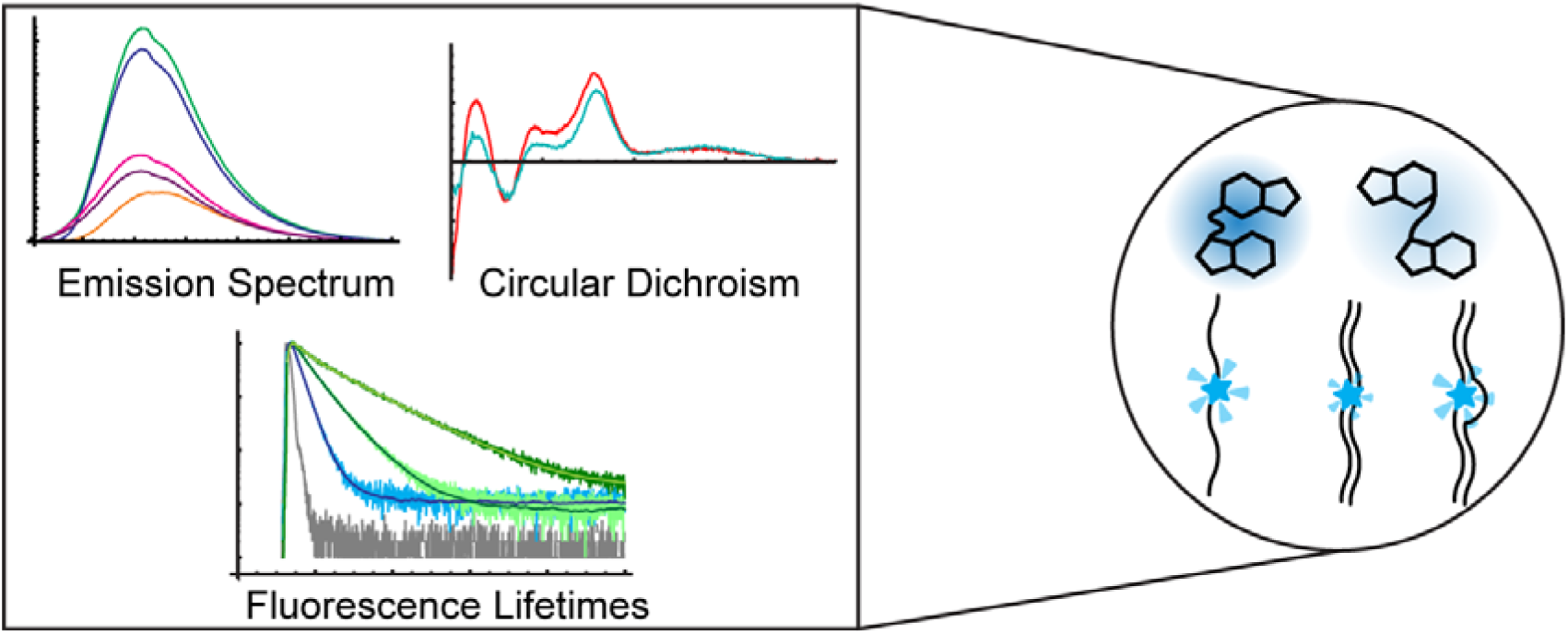

## INTRODUCTION

Fluorescence detection methods are valuable tools for elucidating the structures and dynamics of biopolymers both *in vitro* and *in vivo* (1, 2). For example, fluorescence-based experiments on nucleic acids can be designed to reveal changes in structural features such as base-stacking and hydrogen bonding in response to environmental stimuli (3, 4). Fluctuations in local base-stacking and base-pairing interactions typically occur on a faster timescale than global changes in secondary or tertiary structure, but these local structural reorganizations can be critical for binding interactions and large-scale structural shifts.

The native nucleic acid bases are practically non-fluorescent, with quantum yields on the order of 10^-4^ and excited state lifetimes of less than 1 ps due to rapid, non-radiative de-excitation (5). As a result, external probes must be introduced in order to exploit the structural sensitivity and specificity of fluorescence detection. Fluorescent base analogues (FBAs), such as the widely utilized 2-aminopurine (2-AP), are structural mimics of the native bases with increased fluorescence quantum yields that are designed to maintain the base-pairing properties of their cognate native bases. Certain FBAs, including 2-AP, exhibit environment-sensitive fluorescence properties, which enables their use as probes of local nucleic acid structure (6–9). 2- AP has been extensively characterized and utilized in both DNA and RNA, but it is a structural mimic only of adenine and guanine and experiences a high degree of quenching upon base stacking (6, 10), in some cases rendering highly stacked conformations nearly undetectable (11). To expand the toolbox of probes available to study the ever-expanding list of biologically and medicinally relevant RNA structures, alternative FBAs must be characterized just as thoroughly. Systematic studies of the structure- and sequence-dependence of the fluorescence of any FBA are critical for optimal experimental design and data interpretation.

Pyrrolo-cytosine (pC; Fig. 1A), an analogue of cytosine first synthesized in 2004 (12), has been used sporadically in both DNA (13–15) and RNA (4, 16) and shows promise as a probe of structures and sequences where 2-AP is not optimal. Previous measurements in DNA have shown that pC experiences enhanced fluorescence when flanked by native bases, a property that was attributed to increased solvent shielding in base-stacked conformations (17). It was also observed that base pairing led to fluorescence quenching in pC-labeled DNA oligonucleotides (17), and that base-stacking with guanine opened new nonradiative de-excitation pathways (10, 17, 18). pC therefore shows promise as an alternative FBA that can probe interactions that 2-AP is less sensitive to. However, the mechanisms behind pC’s fluorescence changes upon incorporation into larger nucleic acid structures are poorly understood and have been primarily studied in DNA.

**Figure 1.**
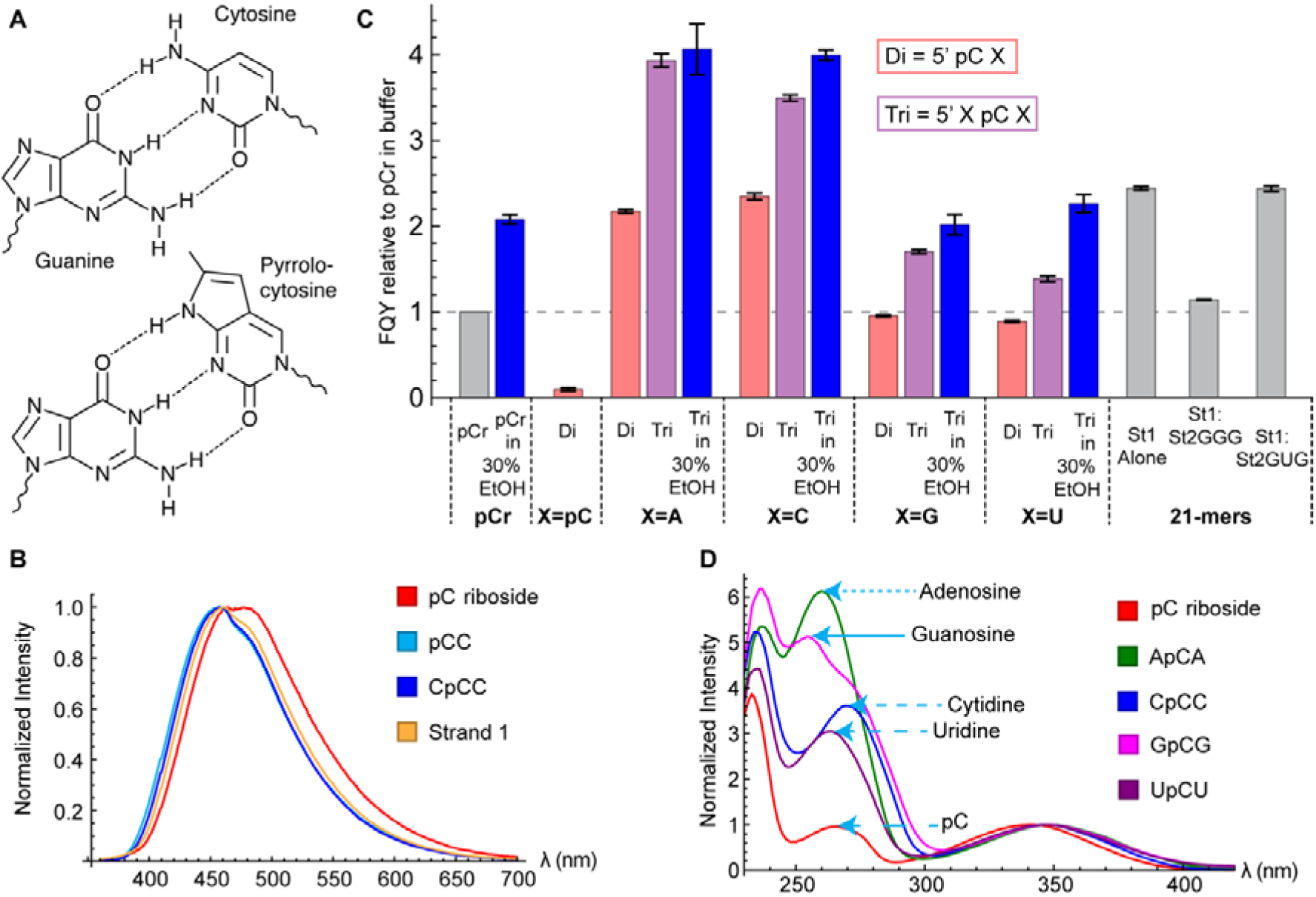
**(A)** Structures of cytosine (top) and pC (bottom) forming Watson-Crick base pairs with guanine. **(B)** Normalized fluorescence emission spectra of pCr (red), pCC (light blue), CpCC (dark blue), and Strand 1 (orange). **(C)** FQY of all oligonucleotides, normalized to value of 1 for pCr in buffer. Error bars show the standard deviation across three separately prepared samples. “Di” indicates a dinucleotide with the base indicated by “X” to the 3’ side of pC. “Tri” indicates a trinucleotide with the base indicated by “X” to the 5’ and 3’ sides of pC. Samples prepared in 30% ethanol are indicated by blue. **(D)** Normalized fluorescence excitation spectra of pCr (red), ApCA (green), CpCC (blue), GpCG (magenta), and UpCU (purple). Approximate peak absorbance wavelengths of each native base (260 nm, 271 nm, 254 nm and 262 nm for A, C, G, and U, respectively) and pC itself (265 nm) are indicated by the tips of the arrows.

In this work, we utilized a variety of spectroscopic methods to probe the impacts of local structure and sequence on pC’s fluorescence, generating a map of its photophysical landscape within RNA that will enable the informed design of probe sequences and deeper interpretation of the data obtained on them. We show that its emissive properties are more complex in RNA than previously observed in DNA and that base-stacked conformations of pC are generally brighter than unstacked conformations, the opposite behavior to 2-AP. This work demonstrates the rich structural information that will be obtainable using pC in biologically relevant RNA contexts, such as ligand binding pockets or regions of fluctuating hybridization status.

## MATERIAL AND METHODS

### Reagents and General Methods

All oligonucleotides were obtained from Dharmacon (Horizon Discovery) and deprotected and desalted by the manufacturer. 21-mers (Table 1) were additionally HPLC-purified by the manufacturer. Stock solutions were prepared in DEPC-treated water and their concentrations were determined by A260 using extinction coefficients provided by the manufacturer. pC riboside (pCr) was obtained from Biosearch Technologies (PYA11092-B010). In most cases, spectroscopic studies were performed with oligonucleotides at a concentration of 5 µM in a buffer consisting of 20 mM sodium phosphate (pH 7.4) and 100 mM NaCl. Where indicated, the buffer additionally included 30% v/v ethanol. All data on pC dinucleotide (pCpC) were recorded at a concentration of 1 µM. Spectra requiring deep UV excitation of 21- mers (absorbance, CD and fluorescence excitation) were recorded at 1 µM to reduce the inner filter effect. Double-stranded complexes were prepared by combining pC-labeled and unlabeled strands in a 1:1.1 ratio (slight excess of unlabeled strand) and annealing them by heating to 90°C for 2 minutes and allowing to cool in air for 10 minutes. Cuvettes were soaked in a solution of 1% Hellmanex at 35°C for 30 minutes and dried using N_2_ before use. Unless otherwise noted, all spectroscopic measurements were done at 20°C.

**Table 1.**
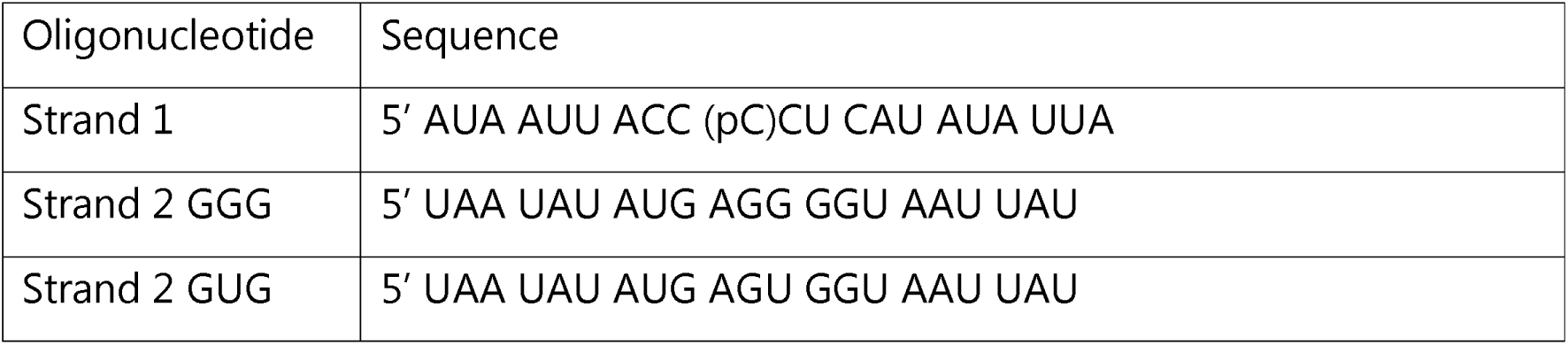
Sequences of the 21-mers used in this study.

### Purification of pCpC

pCpC (Fig. S1A) was obtained as a crude synthesis from Dharmacon and was purified via preparative thin-layer chromatography (TLC). A 20×20 cm 250-micron silica gel TLC plate (Millipore Sigma 1.05715.0001) was first run using pure methanol followed by drying using a gentle flow of N_2_ overnight. Crude pCpC was then deposited in a line along the origin using a gel-loading pipette tip at a density of approximately 1.25 nmol/mm, leaving at least 3 cm empty on each edge (19). Based on established protocols for separating purine derivatives (20), a solvent system composed of *n*-butanol, acetone, 33% ammonia and water (50:40:3:15) was used for separation. The plate was run to a height of 1 cm below the top, dried under gentle N_2_ flow on a hot plate set to 32°C for 30 minutes, and then run a second time to improve separation (19). Bands visible under UV illumination at 254 nm (Fig. S1B, Table S1) were cut out using a razor blade (∼0.25 mL of silica gel) and transferred to 50 mL conical tubes. To extract each band, 1:1:1 water-methanol-acetonitrile was added in a volume based on Eq. 1 (19) and vortexed for 5 minutes before centrifuging at 1500 x *g* for 15 minutes.

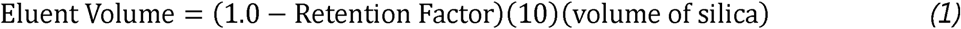

The supernatant was decanted and the extraction was repeated a second time to improve yield. Each extraction’s supernatant was centrifuged three times at 1500 x *g* for 15 minutes and decanted between cycles to remove debris. The supernatant was distributed into 5 mL Eppendorf tubes and dried under vacuum at 45°C. The dry samples were redissolved in DEPC-treated water, filtered using an Ultrafree-MC Centrifugal Filter (0.1 µm pore size), and analyzed via mass spectrometry to identify bands and assess purity (Fig. S1C). Mass spectra were acquired on a Synapt G2-si mass spectrometer (Waters Corp.) in negative ion mode. The samples were sprayed from 3-5 µL of solution at a concentration of 25 µM using nano-Electrospray Ionization (nESI) from a borosilicate capillary with an inner diameter of 0.78 mm pulled to an inner diameter below 1 µm (∼400 nm) and a capillary voltage of 0.3 kV. A mass calibration was performed using Cesium Iodide with a resulting mass accuracy of ∼7 ppm.

### Spectroscopy and Statistical Analyses

#### Steady-State Absorbance and Fluorescence Spectroscopy

Steady-state spectra were collected on an Edinburgh FS5 spectrofluorometer with 1 nm steps and 1 s dwell time. Fluorescence excitation spectra were collected by scanning the excitation monochromator from 220 nm to 420 nm with a bandwidth of 2 nm while monitoring emission at 460 nm with a bandwidth of 3-5 nm, selected to avoid detector saturation. Absorbance spectra were collected in the same manner using the transmission detector on the FS5, with three sequential scans being collected and averaged together. Fluorescence emission spectra were collected by scanning the emission monochromator from 360 nm to 700 nm with a bandwidth of 2 nm while exciting at 340 nm with a bandwidth of 3-5 nm. Fluorescence excitation spectra of 21-mers were corrected for the inner filter effect using the corresponding absorbance spectra and then smoothed with a moving average over four data points. Tables 2 and 3 and Fig. 1C contain relative fluorescence quantum yields (FQYs) of pC and molar absorptivity at 347 nm (ε_347_). Using pCr, which has an FQY of 0.023±0.002 (17), as a reference, the FQY of each oligonucleotide was calculated according to Eq. 2.

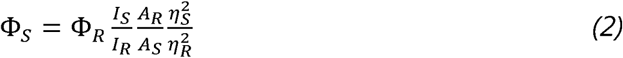

**Table 2.**
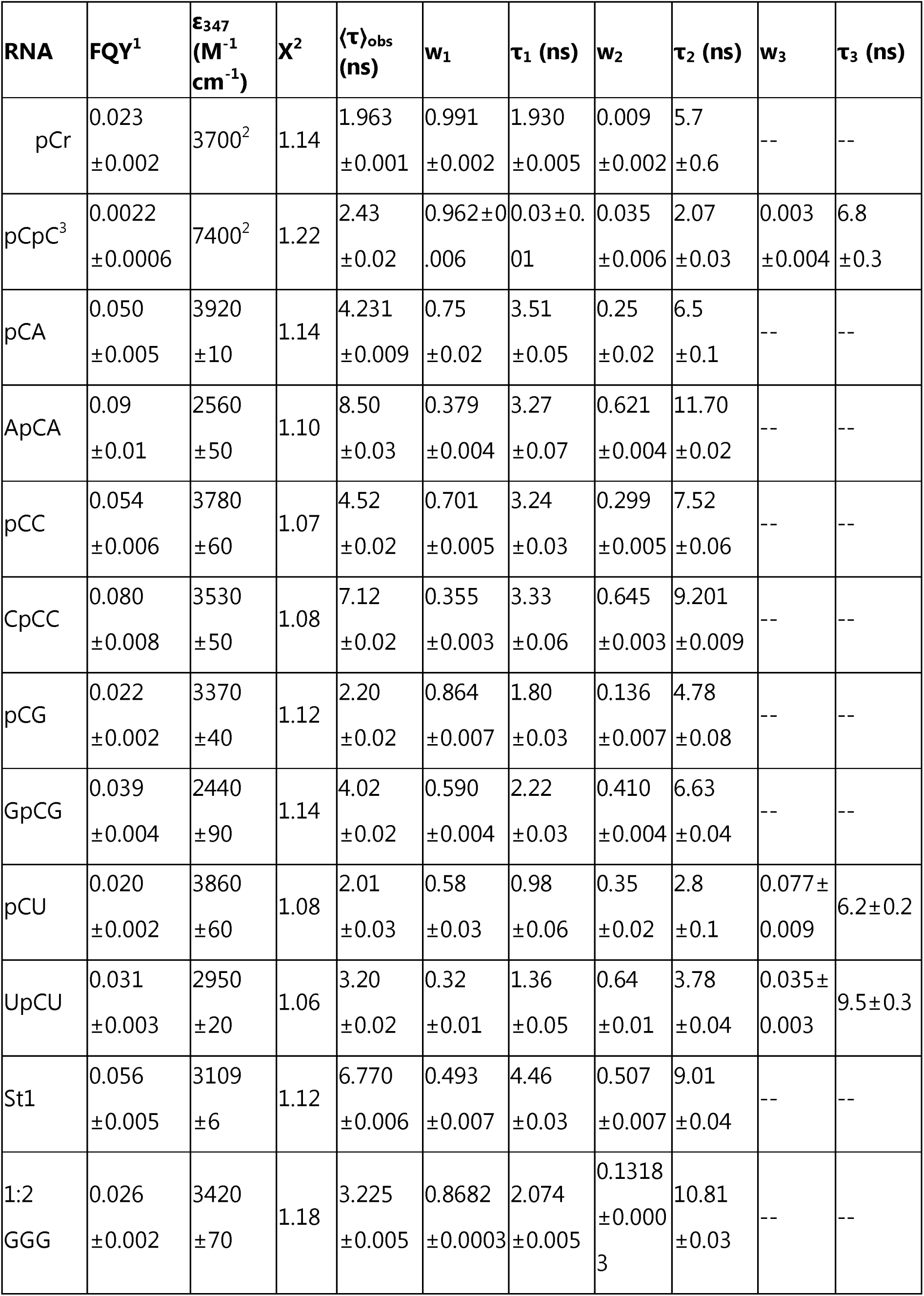
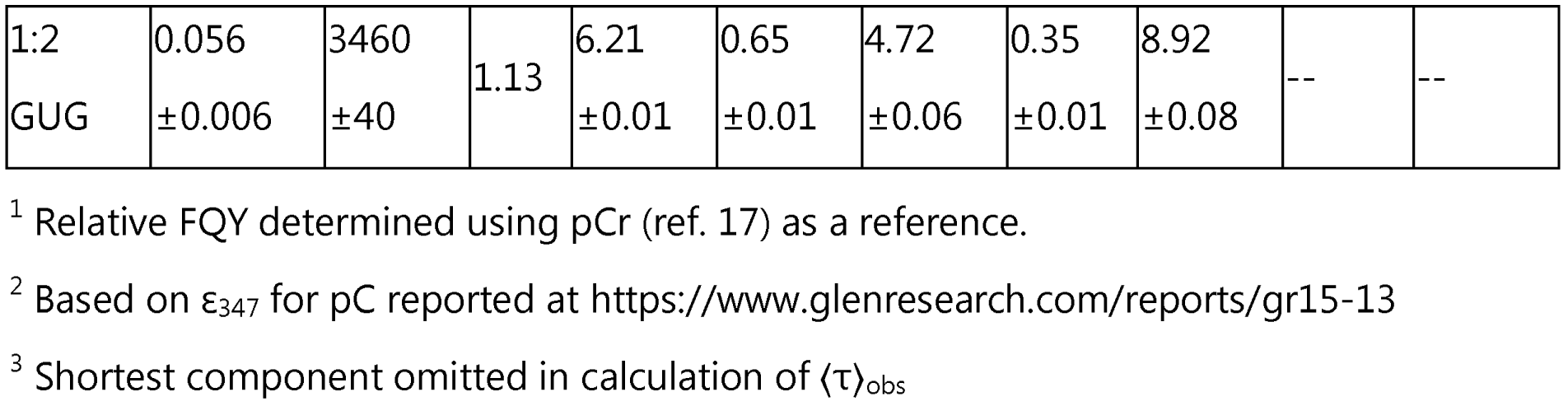
Fluorescence quantum yields (FQY), extinction coefficients (ε_347_) and multi-exponential TCSPC fitting results for all oligonucleotides at 20 °C. Uncertainties for FQY and ε_347_ are the standard deviation across measurements on three samples. Uncertainties for fitting results are the standard deviation across three successive data acquisitions.

**Table 3.** Relative FQY, ε_347_ and bi-exponential TCSPC fitting results for trinucleotides at 20°C in buffer containing 30% v/v ethanol. Uncertainties for FQY and ε_347_ are the standard deviation across measurements on three samples. Uncertainties for fitting results are the standard deviation across three successive data acquisitions.

| RNA | FQY | $\epsilon_{347}$ ($M^{-1}cm^{-1}$) | $\chi^2$ | $\langle\tau\rangle_{obs}$ (ns) | $w_1$ | $\tau_1$ (ns) | $w_2$ | $\tau_2$ (ns) |
| --- | --- | --- | --- | --- | --- | --- | --- | --- |
| pCr | 0.048<br>$\pm 0.005$ | 3990<br>$\pm 70$ | 1.12 | 3.511<br>$\pm 0.004$ | 1.00 | 3.511<br>$\pm 0.004$ | -- | -- |
| ApCA | 0.09<br>$\pm 0.02$ | 2670<br>$\pm 80$ | 1.07 | 8.298<br>$\pm 0.008$ | 0.494<br>$\pm 0.004$ | 4.36<br>$\pm 0.03$ | 0.506<br>$\pm 0.004$ | 12.15<br>$\pm 0.04$ |
| CpCC | 0.092<br>$\pm 0.009$ | 3740 $\pm 20$ | 1.14 | 7.63<br>$\pm 0.02$ | 0.57<br>$\pm 0.02$ | 4.9<br>$\pm 0.1$ | 0.43<br>$\pm 0.02$ | 11.2<br>$\pm 0.2$ |
| GpCG | 0.046<br>$\pm 0.007$ | 2248<br>$\pm 5$ | 1.08 | 4.452<br>$\pm 0.002$ | 0.614<br>$\pm 0.006$ | 2.68<br>$\pm 0.02$ | 0.386<br>$\pm 0.006$ | 7.28<br>$\pm 0.04$ |
| UpCU | 0.052<br>$\pm 0.007$ | 2420<br>$\pm 90$ | 1.14 | 4.06<br>$\pm 0.02$ | 0.68<br>$\pm 0.01$ | 3.00<br>$\pm 0.06$ | 0.32<br>$\pm 0.01$ | 6.28<br>$\pm 0.08$ |

Φ is the fluorescence quantum yield while S and R represent sample and reference, respectively. I is the integrated fluorescence intensity, A is the absorbance at the excitation wavelength, and η is the refractive index of the solvent, which in this work was identical for the samples and reference. ε_347_ was calculated by dividing the absorbance at 347 nm by the concentration, which was determined *in situ* via A260 for single-stranded oligonucleotides. dsRNA samples were assumed to be 5 µM based on the measured concentrations of the ssRNA stocks they were prepared from. The mean and standard deviation of measurements on 3 separately prepared samples are reported.

#### Time-Correlated Single Photon Counting (TCSPC)

TCSPC measurements were performed on an Edinburgh FS5 spectrofluorometer equipped with an EPLED-320 (pulse width of 925.3 ps, wavelength of 325.5 nm, and FWHM bandwidth of 12.5 nm) and a high-speed PMT. Decays were collected to a peak count of 10,000 over a 100 ns window divided into 2048 channels with an LED pulse rate of 10 MHz. Reverse mode, in which the start and stop triggers of the counting mechanism are reversed, was utilized to decrease collection time. Emission was monitored at 460 nm with a bandwidth of 29.9 nm. Instrument response functions (IRFs) were recorded using buffer by tuning the emission monochromator to the LED wavelength. A typical IRF full-width at half maximum (FWHM) was ∼1300 ps. In ideal conditions, time components up to 1/10 this length can be resolved with the time width of each channel (here, 50 ps) setting a lower limit. Data were analyzed in two ways, detailed below: using exponential reconvolution fits in Edinburgh’s instrumental software, Fluoracle, and using lifetime distribution fits in open-source Matlab software (21) developed for use with other FBAs (22). For each sample, three decays were collected sequentially, and each was fitted individually. The mean and standard deviation of the resulting values across the three fits is reported.

#### Exponential Reconvolution Fit Analysis

This approach identifies an exponential function with one or more components that describes the decay of the molecule (Eq. 3), where **τ**_i_ represents the time constant of component *i* and w_i_ represents its population weight.

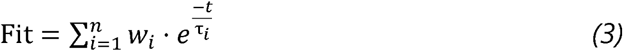

This fitting procedure iterates over time components of an initially proposed fit until it returns an equation that, when combined with the measured IRF, returns the measured decay. The average observed lifetime ⟨**τ**⟩_obs_ is calculated as a weighted average of decay components (Eq. 4).

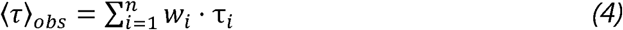

#### Lifetime Distribution Fit Analysis

Lifetime distribution analysis uses a similar approach to exponential reconvolution fit analysis where fitting parameters are iterated until the fit returned, when combined with the IRF, matches the measured decay. Here, we use a previously published model in which the fitting equation allows both exponential decay components and components described by gamma probability distributions, where ɑ_i_ and *κ*_i_ are the shape and scale parameters of the gamma distribution, respectively (21, 22; Eq. 5).

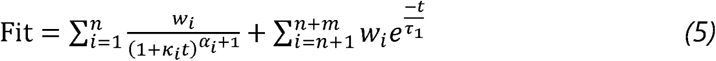

If a decay component is best described by a gamma distribution, the mean of the distribution (Eq. 6) is taken as its time constant.

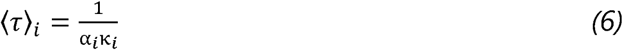

⟨**τ**⟩_obs_ is calculated as above using ⟨**τ**⟩_i_ in place of **τ**_i_ for any gamma components.

#### Selection of Fluorescence Decay Models

The quality of a fit is determined by two parameters: 1) residuals and 2) reduced χ^2^. Residuals with a magnitude less than 5 and a random distribution combined with a reduced χ^2^ of 1-1.3 are indicative of an acceptable fit (21). Mono-exponential fits were initially tested for all decays and additional exponential or gamma components were tested until the fit was acceptable. In cases where the decay is equally well fit by more than one model, the fit with the fewest number of parameters is selected.

#### Circular Dichroism and Fluorescence-Detected Circular Dichroism

CD and FDCD spectra were recorded as previously described for 2-AP (11) with minor modifications detailed below. Analysis was performed in Mathematica (Wolfram Alpha). Spectra were recorded on a Jasco J-1500 CD spectrometer equipped with a FDCD-551 attachment. A long-pass colored glass filter with a cut-on wavelength of 420 nm was placed in front of the detector. The measurement channels “FDCD” and “DC” were acquired with the gain on the detector selected such that the maximum DC signal was approximately 1 V. Scans were performed in continuous scanning mode from 200 to 410 nm with a 4 nm excitation bandwidth, a 4 s integration time, and a scan speed of 20 nm/minute. Due to its low fluorescence intensity, pCpC was collected in step-scan mode from 200 to 410 nm with a 4 nm excitation bandwidth, a 4 s integration time, and a data pitch of 1 nm. Multiple scans were acquired, ranging from 5 for ApCA to 20 for pCpC, and averaged together to obtain adequate signal-to-noise ratio (SNR). CD spectra were collected using the same scan parameters as FDCD, though fewer cycles were typically needed to achieve adequate SNR, and were background-corrected using buffer scans. Sodium fluorescein was used as an achiral reference fluorophore to background-correct all FDCD spectra (23). For each measurement of a pC-containing sample, scans were also performed on buffer, water and an aqueous fluorescein solution at a concentration yielding a maximum DC signal of approximately 1 V at a similar detector gain setting to that used for the pC-containing sample (the required fluorescein concentrations ranged from 0.1-3 nM). To correct for stray or scattered excitation light, the FDCD and DC channels of the buffer scan were subtracted from the corresponding channels of the pC scan, and the FDCD and DC channels of the water scan were subtracted from the corresponding channels of the fluorescein scan. The quantity θ_F_ = FDCD/DC was then computed for both pC and fluorescein. θ_F_ for fluorescein was smoothed using a Wiener filter and then subtracted from θ_F_ for the pC-containing sample.

Established theory (24) was used to convert the background-corrected θ_F_ into more readily interpretable quantities. One such quantity is the dissymmetry factor g_F_=Δε_F_/ε_F_, where ε_F_ is the extinction coefficient of the fluorescent species in the sample and Δε_F_ is its difference in extinction coefficient under excitation with left- and right-handed circularly polarized light (ε_L_-ε_R_). g_F_ was calculated using Eq. 7 with θ_F_ and the CD (in millidegrees) and absorbance (A, in absorbance units) spectra of the sample as inputs:

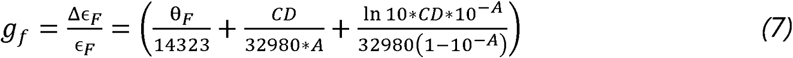

The analogous quantity from standard CD spectra was calculated as g = CD/(32980*A).

CD spectra are most commonly displayed as Δε, and obtaining an analogous quantity from FDCD, Δε_F_, requires an ε_F_ spectrum to multiply through Eq. 7. When multiple species and energy transfer (ET) are present, ε_F_ must be replaced with a population-, quantum yield- and ET efficiency-weighted sum over all species (24):

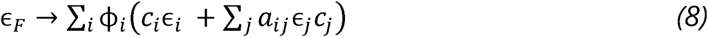

where *i* counts over all fluorescent species, *j* counts over all the species that can transfer energy to fluorescent species *i*, *Φ* _i_ is the fluorescence quantum yield of species *i*, *a* _ji_ is the efficiency of ET from species *j* to species *i*, *ε*_i_ and *ε* _j_ are the extinction coefficients of species *i* and ET donor *j*, respectively, and *c*_i_ and *c*_j_ are their concentrations. As previously noted (23), this quantity is reasonably approximated by the fluorescence excitation spectrum. To obtain fluorescence excitation spectra across the entire wavelength range probed by FDCD (200-420 nm), the DC signal at each wavelength was divided by the lamp intensity (measured with a power meter, Thorlabs S120VC) and the absorbance spectrum used to correct the DC spectrum for the inner filter effect. Fluorescence excitation spectra computed in this manner agreed well with spectra recorded on the FS5 over its accessible wavelength range (>230 nm). The excitation spectrum of each sample was converted to units of L/mol*cm by scaling it uniformly such that ε_347_ had the value listed in Table 2. g_F_ was multiplied by this spectrum in order to obtain Δε_F_.

### Melting Curves

Melting curves were recorded on a Jasco J-1500 CD spectrometer in a low headspace, 1 cm pathlength cuvette (Starna 26.160/LHS). The samples were heated at 1°C/min from 4°C to 80°C and absorbance was monitored at 260 nm. T_M_ values were extracted by fitting the melting curves in Mathematica, as previously described (3).

### Computational Methods

Based on the work of Nguyen et al. (18), the ground state molecular geometries of G, C, and pCr nucleobases were optimized at the MP2/6-31G(d,p) level. These optimized monomers were combined into the trimers GpCG and CpCC in an A-form helix, the geometry of which was obtained from the literature (10). The vertical excitation energies for the first ten excited states were calculated using the configuration interaction singles with perturbative second order corrections (CIS(D)) at the aug-cc-pVDZ (25) level of theory, selected based on computational recommendations for pC-containing systems from Nguyen et al (18). Following excited state calculations, the natural transition orbitals (NTOs) (26) were calculated for each excited state at the same level of theory. All calculations were carried out in Gaussian09 (27).

## RESULTS

We first investigated the impact of base-stacking and sequence context on the fluorescence of pC by comparing the steady-state and time-resolved emission properties of di- and trinucleotides containing pC adjacent to each native RNA base. Each dinucleotide studied consisted of pC to the 5’ side of a native base, and each trinucleotide consisted of pC flanked by two of the same native base. The fluorescence emission spectra of all dinucleotides exhibit a blue-shift in peak wavelength along with a change in lineshape relative to pCr (Fig. 1B). Minimal further changes in the spectrum are observed in the corresponding trinucleotides (Fig. S2). This hypsochromic shift in emission wavelength and suppression of a long-wavelength (red) vibronic shoulder is consistent across all sequence contexts, indicating that local changes in the solvation environment impact pC’s emission spectrum regardless of the identity of the neighboring native base. When adjacent to one A or C residue, the raw fluorescence intensity (Fig. S3) and FQY (Fig. 1C; Table 2) of pC approximately double relative to pCr and increase further in the corresponding trinucleotides. In contrast, the presence of one adjacent G or U residue leaves pC’s FQY nearly unchanged, while it is enhanced by about 50% in the corresponding trinucleotides.

Absorbance and fluorescence excitation spectra of the di- and trinucleotides all show the expected ∼345 nm feature resulting from direct excitation of pC, but that feature’s peak wavelength is slightly red-shifted and its extinction coefficient is lower in the trinucleotides relative to pCr (Fig. S3; Table 2). In fluorescence excitation spectra, significant differences in relative intensity and lineshape are observed in the 250-300 nm region (Fig. 1D). This could be attributed to energy transfer from the native bases such that excitation of an adjacent native base results in pC fluorescence. Indeed, the peak wavelength of this feature closely matches the known absorption maximum of the native base present in each oligonucleotide (Fig. 1D) (28). When normalizing by the 345 nm peak to account for differences in overall emission intensity, the variation in the intensity of the excitation peak in the 260 nm region suggests a moderately sequence-dependent energy transfer efficiency. The intensity is highest in pCA and ApCA, which could be explained by the fact that A has the largest extinction coefficient (15,400 M^-1^cm^-1^) of all four nucleoside monophosphates (NMPs). However, the peak intensity near 260 nm is almost identical in pCC and pCG despite CMP having an extinction coefficient of ∼9,000 M^-1^cm^-1^ compared to GMP’s 13,700 M^-1^cm^-1^, and is substantially lower in pCU despite UMP having an extinction coefficient (10,000 M^-1^cm^-1^) close to that of CMP (28). This suggests that energy transfer from G or U is less efficient than from A and C, and/or that the most highly fluorescent conformations of pCG or pCU have low energy transfer efficiencies. In trinucleotides (Fig. 1D), the intensity of the ∼260 nm feature in the excitation spectrum more closely tracks with extinction coefficient (A>G>>C), with the exception that it is again lower in UpCU than would be predicted by the extinction coefficient of UMP.

To probe the distribution of structures present in solution, we determined the excited state lifetimes of the oligonucleotides under study using TCSPC (Fig. 2). The resulting fluorescence decays were fitted with exponential or distribution models containing different numbers of decay components (see methods for details). Typically, each decay component is interpreted as a distinct conformational state or a distribution of similar or rapidly interconverting states, with its weight reflecting the prevalence of that state (21, 22). All pC-containing oligonucleotides have longer lifetimes than pCr and their decays exhibit evidence of conformational heterogeneity (Fig. 2). Specifically, the decays of all di- and trinucleotides require at least two components to be adequately fit (χ^2^=1.06-1.14), which likely originate from a combination of base-stacked and unstacked states (Table 2). Unlike oligonucleotides containing A, C, and G, the decays of pCU and UpCU are significantly better fit with either a tri-exponential (“3E”) model (χ^2^=1.08 for pCU; 1.06 for UpCU) or a model containing a fast exponential component and a slower gamma distribution component (“1E+1G”, χ^2^=1.08 for pCU; 1.09 for UpCU) than with the simple bi-exponential (“2E”) model that is adequate for all the others (χ^2^=1.25 for pCU; 1.22 for UpCU). More detail about comparison of fitting models is given in Supplementary Note 1.

**Figure 2.**
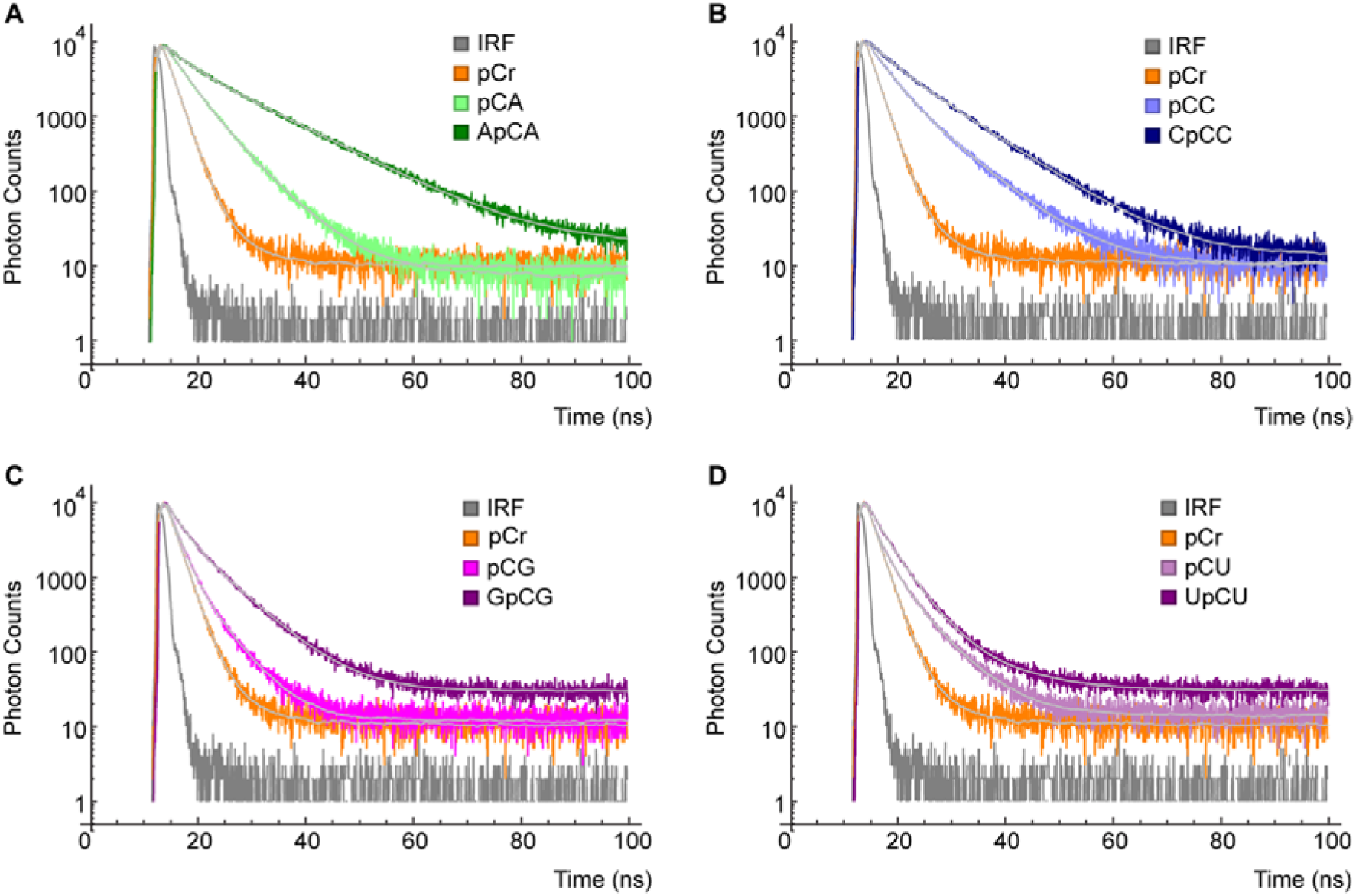
TCSPC measurements of pCr and di- and trinucleotides. **(A)** pCA and ApCA, **(B)** pCC and CpCC, **(C)** pCG and GpCG, **(D)** pCU and UpCU. Bi-exponential **(A, B, C)** or tri-exponential **(D)** fits are overlaid in light gray. A typical IRF is plotted in gray.

The shortest decay component is dominant in all dinucleotides, while the population distribution shifts away from the fastest decay component in trinucleotides (Table 2). In addition, the lifetime associated with the faster component is nearly unchanged between A-, C- and G-containing dinucleotides and their corresponding trinucleotides, while the slower component lengthens significantly in the trinucleotides. We hypothesized that the longer decay component describes a base-stacked conformational state, as this state would be more protected from solvent in a trinucleotide and would be absent in pCr. 3E fits to the decays of pCU and UpCU show that the trinucleotide exhibits longer time constants in all decay components and a larger population associated with the intermediate τ∼3 ns component, while the population of the longest decay component remains low (<10%) in both. Though pCU and UpCU sample a more complex distribution of conformational states, we hypothesized that, like the other di- and trinucleotides, the longer of their two heavily populated components describes a base-stacked structure.

To confirm the identities of these conformational states, we used differences in solvent polarity to alter the base-stacking equilibrium. Ethanol has previously been shown to enhance the raw fluorescence intensity of pCr (8) and shift the base-stacking equilibrium of di- and trinucleotides to favor the unstacked state (11, 29). We performed measurements on pCr and all four trinucleotides in the presence of 30% (v/v) EtOH, hypothesizing that ethanol would induce an increase in the population weights of decay components originating from unstacked conformations. In 30% EtOH, an increase in FQY is observed for pCr, CpCC, GpCG and UpCU (Table 3; Fig. 1C), and both decay time constants of all four trinucleotides lengthen (Table 3; Fig. 3; Fig. S4), attributed to the enhancement of fluorescence in a less polar environment (8). Notably, CpCC and ApCA show a significant increase in the population weight of the shorter decay component in 30% EtOH and a compensatory decrease in the population weight of the longer one (Table 3). The same shift occurs to a lesser degree in GpCG. This shift in population weight supports the notion that the longer decay component originates from a base-stacked conformation and the shorter decay component, which is favored in less polar solvent, originates from an unstacked conformation. Unlike in buffer, the decay of UpCU in 30% EtOH is fit well (χ^2^=1.14) with a 2E model that lacks the low population, slowly decaying component observed in buffer. EtOH impacts the two shorter components in the same manner as observed in the other trinucleotides, with their time constants lengthening and population weight shifting to the shortest decay component. Therefore, while adjacent uracil residues add complexity to the fluorescence decays of pC, the observed population shift suggests a similar structural interpretation.

**Figure 3.**
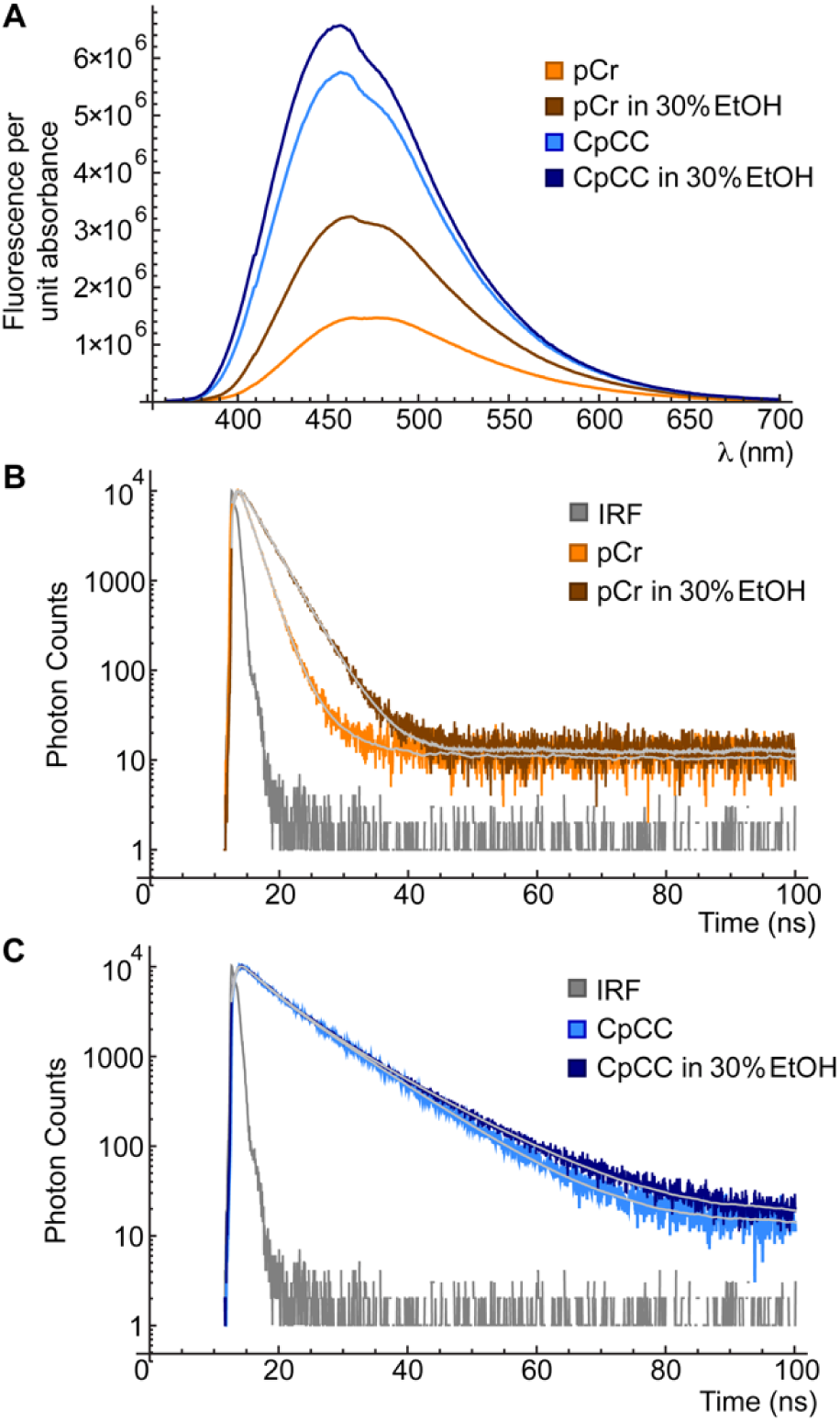
**(A)** Fluorescence per unit absorbance of pCr (orange) and CpCC (blue) in buffer (lighter shades) or buffer + 30% EtOH (darker shades). **(B)** TCSPC of pCr in buffer or buffer + 30% EtOH. **(C)** TCSPC of CpCC in buffer or buffer + 30% EtOH. Multi-exponential fits are overlaid in light gray (mono-exponential for pCr in 30% EtOH and bi-exponential for all others).

We further investigated the relationship between base-stacking, fluorescence quantum yield and energy transfer by comparing the oligonucleotides’ circular dichroism (CD) and fluorescence-detected circular dichroism (FDCD) spectra (Fig. 4). FDCD spectra reveal the CD signals of fluorescent species in a sample, weighted by their populations and fluorescence quantum yields, by measuring the difference in emission intensity of a sample under excitation with left-handed and right-handed circularly polarized light (LCP and RCP, respectively) (23). A large FDCD signal indicates that the brightest species in a sample have large CD signals (typically associated with stacked structures), while a small FDCD signal indicates that the brightest species have small CD signals. Raw FDCD spectra were converted (details in methods) into the quantity Δε_F_, which is analogous to the quantity Δε commonly reported in standard CD (Fig. 4). Δε_F_ spectra will henceforth be referred to as “processed FDCD spectra”. We also considered the dissymmetry factor g_F_=Δε_F_/ε_F_, analogous to g=Δε/ε obtained from standard CD, which can be computed directly from raw data (details in Supplementary Note 2 and Fig. S5) (24).

**Figure 4.**
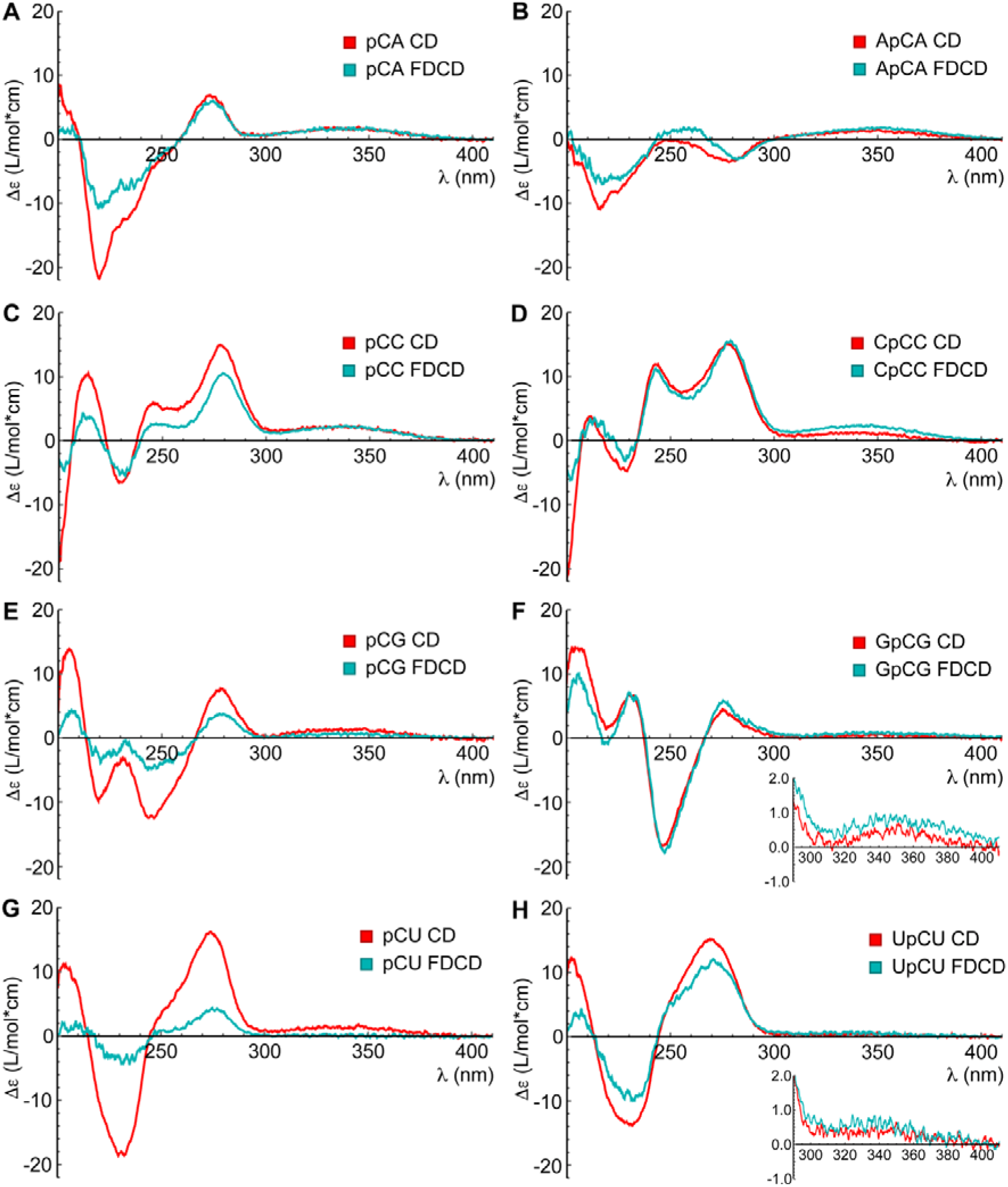
CD (red) and processed FDCD (cyan) spectra of di- and trinucleotides. **(A)** Spectra of pCA. **(B)** ApCA. **(C)** pCC. **(D)** CpCC. **(E)** pCG. **(F)** GpCG (inset: close-up of long-wavelength region). **(G)** pCU. **(H)** UpCU (inset: close-up of long-wavelength region).

In the dinucleotide with the highest FQY, pCC, the CD and processed FDCD spectra agree roughly in lineshape and intensity, with the agreement being nearly quantitative in the long-wavelength band that originates from direct absorption by pC (Fig. 4). In CpCC, the FDCD and CD lineshapes are again very similar, but the longest-wavelength band is more intense in the FDCD spectrum. In both pCC and CpCC, the similarity between the FDCD and CD spectra below 300 nm is strong evidence of the energy transfer that was surmised from fluorescence excitation spectra, showing that differential absorption of LCP and RCP by adjacent C residues is efficiently transduced into differential emission by pC. Together, this indicates that the bright conformations of CpCC have large CD signals >300 nm and efficient energy transfer, both of which would be expected for stacked conformations. In contrast, the FDCD spectrum of pCU is far weaker than its CD spectrum across the entire spectral range of 200-400 nm. Its FDCD and CD lineshapes are similar below 300 nm, indicating that the majority of this signal again originates from differential absorption of LCP and RCP by U followed by energy transfer. The weak FDCD signal indicates that the bright conformations of pCU are those that have weak CD signals, and thus are likely unstacked, consistent with the fact that pCU is slightly quenched relative to pCr (Fig. 1C). In UpCU, the intensity of the FDCD spectrum approaches that of the CD spectrum, showing that stacked conformations are now bright, which is consistent with the increase in FQY of UpCU relative to pCr. Di- and trinucleotides containing A show very similar patterns of behavior to those containing C, and G exhibits similar patterns of behavior to U (Fig. 4).

Numerous studies utilizing 2-AP have placed probes at two adjacent sites (30–35) such that a base-stacked structure forms a classic “H-type” exciton-coupled homodimer in which the lowest-energy excited state is dark (30, 36). We investigated whether a pC homodimer would exhibit similar properties, given the competing factor that base-stacking in some cases enhances the fluorescence of pC. We found that the dinucleotide pCpC was quenched approximately 10-fold compared to pCr, greatly exceeding any pC monomer-containing oligonucleotide (Table 2). As seen in 2-AP dinucleotide (11, 29), pCpC exhibits a strong sigmoidal feature in its CD spectrum at short wavelength and an FDCD spectrum that is far weaker (Fig. S6). This suggests that the base-stacked conformations that give rise to large CD signals are strongly quenched, and thus contribute minimally to the FDCD signal. Furthermore, in contrast to all pC monomer-containing oligonucleotides, fitting the fluorescence decays of pCpC required a time component too short to be accurately resolved with our benchtop TCSPC instrument (Table 2). Together, these results suggest that pC’s distinctive emissive properties may be best exploited using single labels, as the nuanced dependence of its fluorescence on local structure is obscured by the strong self-quenching that occurs in a homodimer.

To further investigate the mechanism of pC’s observed sequence-dependent fluorescence enhancement, we calculated excitation energies, oscillator strengths and natural transition orbitals (NTOs) at the CIS(D)/aug-cc-pVDZ level of theory for CpCC and GpCG (representing strongly and weakly fluorescence-enhancing native bases, respectively) and their constituent nucleobases (Table 4; Fig. S7). Each trinucleotide was modeled by arranging geometry-optimized monomers in an A-form helical geometry with coordinates obtained from the literature (10). The lowest-energy transition in both trimers is a π□π* transition localized on the pC residue (Fig. S7), with a decreased oscillator strength as compared to the equivalent transition in pC base (34% lower for CpCC and 39% lower for GpCG; Table 4). The S_0_ □ S_1_ excitation energy decreases relative to pC base in both trimers (by ∼0.1 eV in CpCC vs. ∼0.05 eV in GpCG). These results are directionally consistent with the experimentally observed reductions in extinction coefficient (Table 2) and redshifts of the longest-wavelength absorbance band (Fig. S3) in the trinucleotides relative to pCr.

**Table 4.** Summary of excited state calculations for CpCC and GpCG with the S_1_ state of pC base for comparison. Charge transfer states are indicated as “CT”.

| Model | Excited State | Excitation Energy (eV) | Wavelength (nm) | Oscillator Strength | Character of Transition |
| --- | --- | --- | --- | --- | --- |
| pC base | S<sub>1</sub> | 3.77 | 329 | 0.2555 | $\pi \rightarrow \pi^*$ |
| CpCC | S<sub>1</sub> | 3.66 | 338 | 0.1689 | $\pi \rightarrow \pi^*$<br>Localized on pC |
|  | S<sub>2</sub> | 4.21 | 294 | 0.0009 | Localized on pC |
|  | S<sub>3</sub> | 4.53 | 273 | 0.0079 | Localized on pC |
| | S<sub>4</sub> | 4.59 | 269 | 0.0954 | CT pC → 3' C |
|  | S<sub>5</sub> | 4.66 | 266 | 0.1480 | Localized on 5' C |
| | S<sub>6</sub> | 4.67 | 265 | 0.0013 | CT pC → 3' C |
| | S<sub>7</sub> | 4.85 | 255 | 0.0151 | CT pC → 3' C |
|  | S<sub>8</sub> | 5.10 | 243 | 0.0046 | Localized on pC |
|  | S<sub>9</sub> | 5.17 | 240 | 0.0147 | Localized on 5' C |
|  | S<sub>10</sub> | 5.27 | 235 | 0.0029 | Delocalized |
| GpCG | S<sub>1</sub> | 3.70 | 335 | 0.1572 | $\pi \rightarrow \pi^*$<br>Localized on pC |
|  | S<sub>2</sub> | 4.32 | 287 | 0.0062 | Localized on pC |
|  | S<sub>3</sub> | 4.72 | 263 | 0.0049 | Localized on pC |
|  | S<sub>4</sub> | 4.77 | 260 | 0.0159 | Localized on 3' G |
|  | S<sub>5</sub> | 4.79 | 259 | 0.0014 | Localized on 5' G |
|  | S<sub>6</sub> | 4.83 | 257 | 0.0106 | Localized on 5' G |
| | S<sub>7</sub> | 4.90 | 253 | 0.0059 | CT pC → 5' G |
|  | S<sub>8</sub> | 4.97 | 249 | 0.0886 | Localized on 3' G |
|  | S<sub>9</sub> | 5.02 | 247 | 0.1477 | Localized on 5' G |
| | S<sub>10</sub> | 5.04 | 246 | 0.0574 | CT pC → 3' G |

Slight differences in behavior due adjacent native base identity are also observed in higher-energy excited states. The S_4_ and S_5_ states of CpCC show oscillator strengths close to that of S_1_ while both being approximately 1 eV greater in energy. The S_4_ state shows full charge transfer from the pC base to the 3’ C, while the S_5_ state shows localized excitation of the 5’ C (Fig. S7). Likewise, the S_8_ and S_9_ states of GpCG lie approximately 1 eV higher in energy than S_1_ and exhibit high oscillator strengths and excitations localized on the 3’ and 5’ native base, respectively. Unlike the S_4_ state of CpCC, the S_8_ state of GpCG lacks charge-transfer character. However, its S_10_ state is only slightly higher in energy (∼0.01 eV), has an oscillator strength ∼1/3 that of S_1_ state, and does show charge transfer from pC to the 3’ G. Excitation into these native-base-localized states followed by energy transfer to pC is likely responsible for the ∼260 nm features observed in fluorescence excitation and FDCD spectra. With large energy gaps between the bright S_1_ state and the other excited states, GpCG and CpCC lack pathways to quenching such as the low-lying charge-transfer and dark states that have been observed in computational studies of other FBAs such as 2-AP (37) and thienoguanosine (^th^G) (38).

To investigate pC’s potential to report on aspects of RNA structure other than base-stacking, we compared a single-stranded (ss) 21-nucleotide RNA with a central CpCC sequence to a fully complementary double-stranded (ds) RNA containing a G opposite pC and a duplex containing a U opposite pC (mismatch; mm) (Table 1). Coupling steady-state and time-resolved spectroscopic methods again proved to be useful as they are sensitive to different aspects of pC’s local environment. As previously observed in DNA and RNA (4, 17), pC’s fluorescence is quenched significantly when opposite a complementary G (Fig. 1C; Table 2). This is accompanied by the appearance of a dominant (>80% population weight) ∼2 ns component in its fluorescence decay (Table 2). In contrast, mm dsRNA recovers nearly the same fluorescence intensity as ssRNA with comparable decay time constants and weights when analyzed with a bi-exponential model (χ^2^=1.12 for ss; 1.13 for mm). Interestingly, the decay of mm dsRNA is also well described (χ^2^= 1.11) by a single broad distribution of conformations and associated lifetimes (Table 5). This “1G” model fits the decay of ssRNA only moderately well (χ^2^= 1.23) and fits the decay of fully complementary dsRNA very poorly (χ^2^>2), indicating that mm dsRNA can access a more continuous range of conformational states. While base-paired structures are strongly distinguished from unpaired (ss or mm) structures by their fluorescence intensity and decays, we found that single-stranded environments are clearly distinguished from ds or mm by their excitation and emission spectra (Fig. 5). Specifically, fully complementary and mismatched duplexes both show evidence of inter-strand energy transfer in their fluorescence excitation spectra through a blueshift of the ∼260 nm feature toward the absorbance maximum of G. Their emission spectra are blueshifted relative to ssRNA and have a less prominent red vibronic shoulder (Fig. 5).

**Figure 5.**
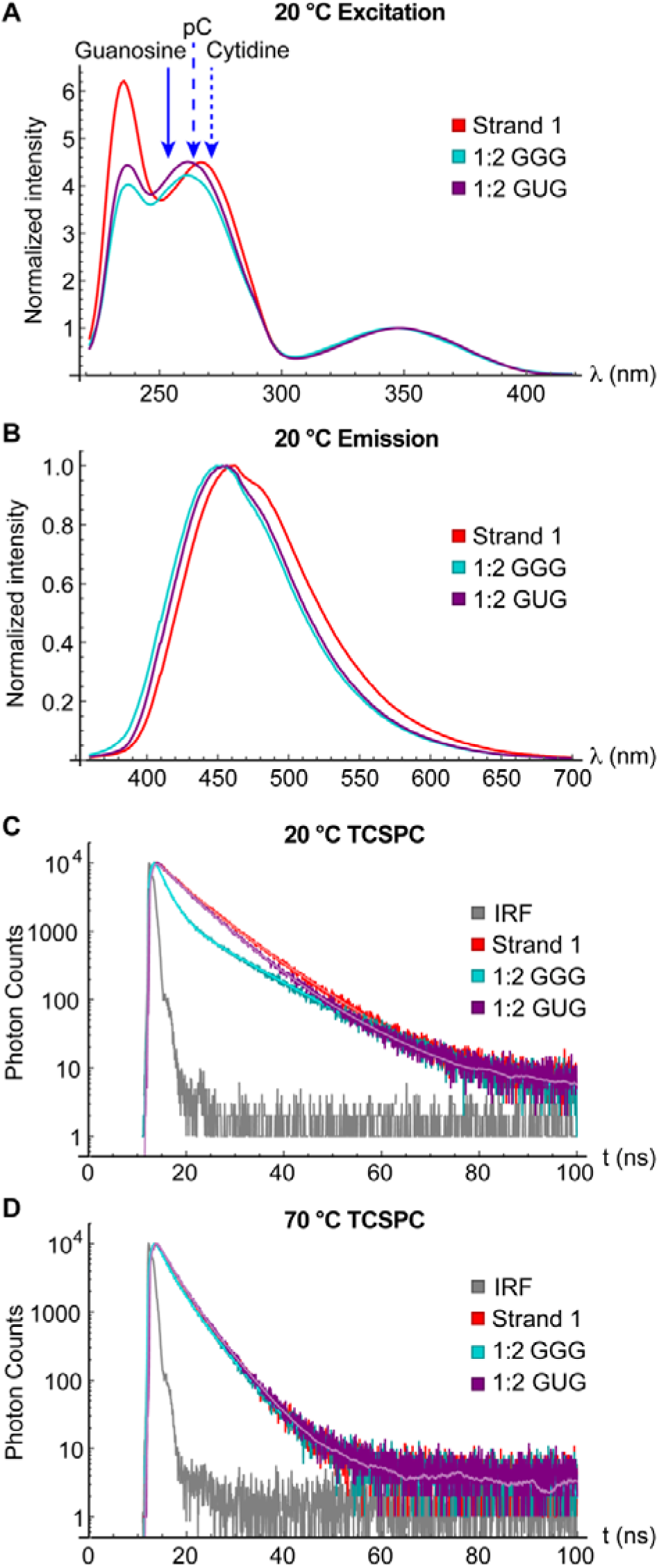
Steady-state fluorescence spectra and TCSPC of single-stranded (“Strand 1”; red), double-stranded (“1:2 GGG”; cyan) and mismatched (“1:2 GUG”; purple) 21-mers. **(A)** Normalized fluorescence excitation spectra of ss, ds and mm RNA at 20 °C. Emission was detected at 460 nm. The approximate peak absorption wavelengths of guanosine (solid blue arrow), cytidine (dotted arrow) and a secondary absorption peak of pC (dashed arrow) are indicated. **(B)** Normalized fluorescence emission spectra of ss, ds and mm RNA at 20 °C under 340 nm excitation. **(C)** IRF (gray) and fluorescence decays of ss, ds and mm RNA at 20 °C. Fits (bi-exponential for Strand 1 and 1:2 GGG; 1 gamma for 1:2 GUG) are overlaid in lighter color. **(D)** IRF (gray) and fluorescence decays of ss, ds and mm RNA at 70 °C. 1 exponential +1 gamma fits are overlaid in lighter color.

**Table 5.**
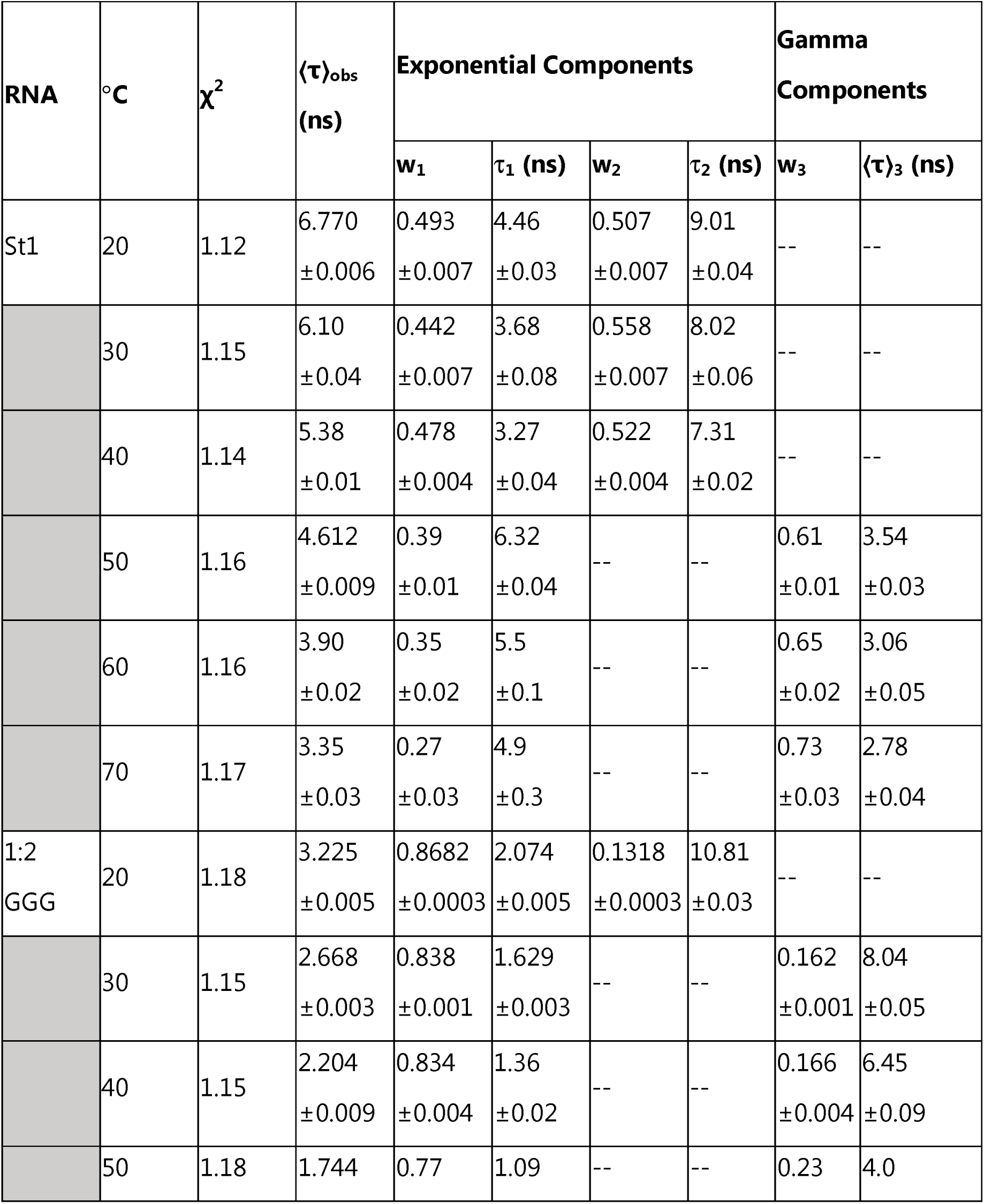
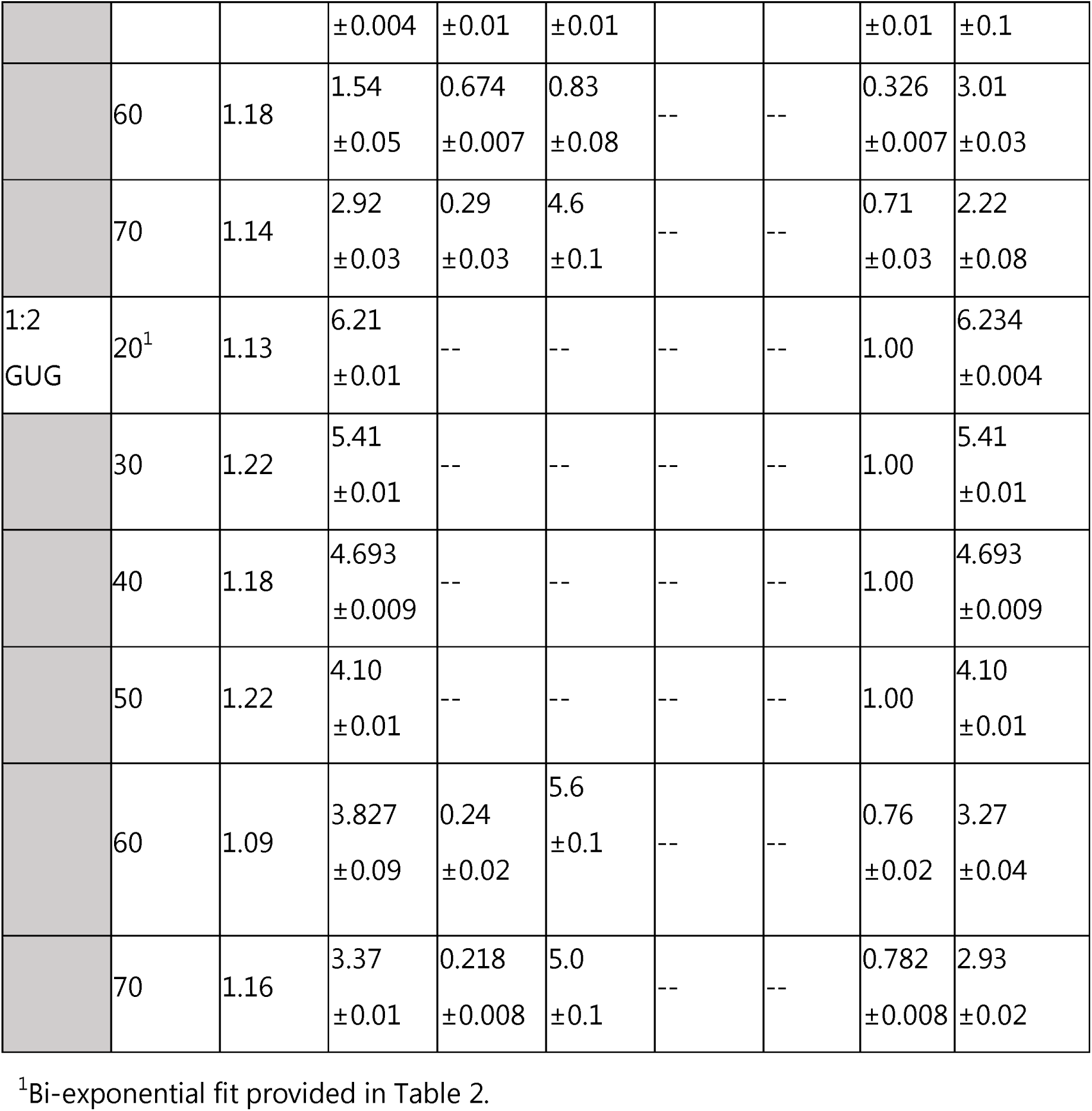
TCSPC fitting results for ss, ds, and mm 21-mers. The optimal fitting model for each circumstance (bi-exponential, 1 exponential+1 gamma, or 1 gamma) is indicated by which columns are occupied. Uncertainties are the standard deviation across three successive data acquisitions.

Unsurprisingly, the CD spectra of fully complementary and mismatched dsRNA are very similar, while the CD spectrum of ssRNA is weaker and has a different lineshape (Fig. 6). The processed FDCD spectra are far weaker than the CD spectra in all cases, which is unsurprising given that they arise only from excitation of pC and bases that transfer energy to it. A positive CD signal is observed at long wavelength in ds and mm RNA, as seen in all di- and trinucleotides (Fig. 4). Notably, the FDCD signal of fully complementary dsRNA is negative at long wavelength, while the long-wavelength features of ss and mm RNAs are positive. This agrees with the observation via TCSPC that pC experiences similar local environments in ss and mm 21-mers, sampling states with lifetimes of approximately 4.5 and 9 ns in both structural contexts when analyzed with a bi-exponential model (Table 2). This stands in contrast to the distinctive (τ=2.1 ns) emissive state of pC in dsRNA. The FDCD signal in the region around 260 nm primarily reflects the optical activity of the native bases that transfer energy to pC (as seen in di- and trinucleotides), and these are observed to adopt a similar geometry in both fully complementary and mismatched duplexes (Fig. 6B).

**Fig 6.**
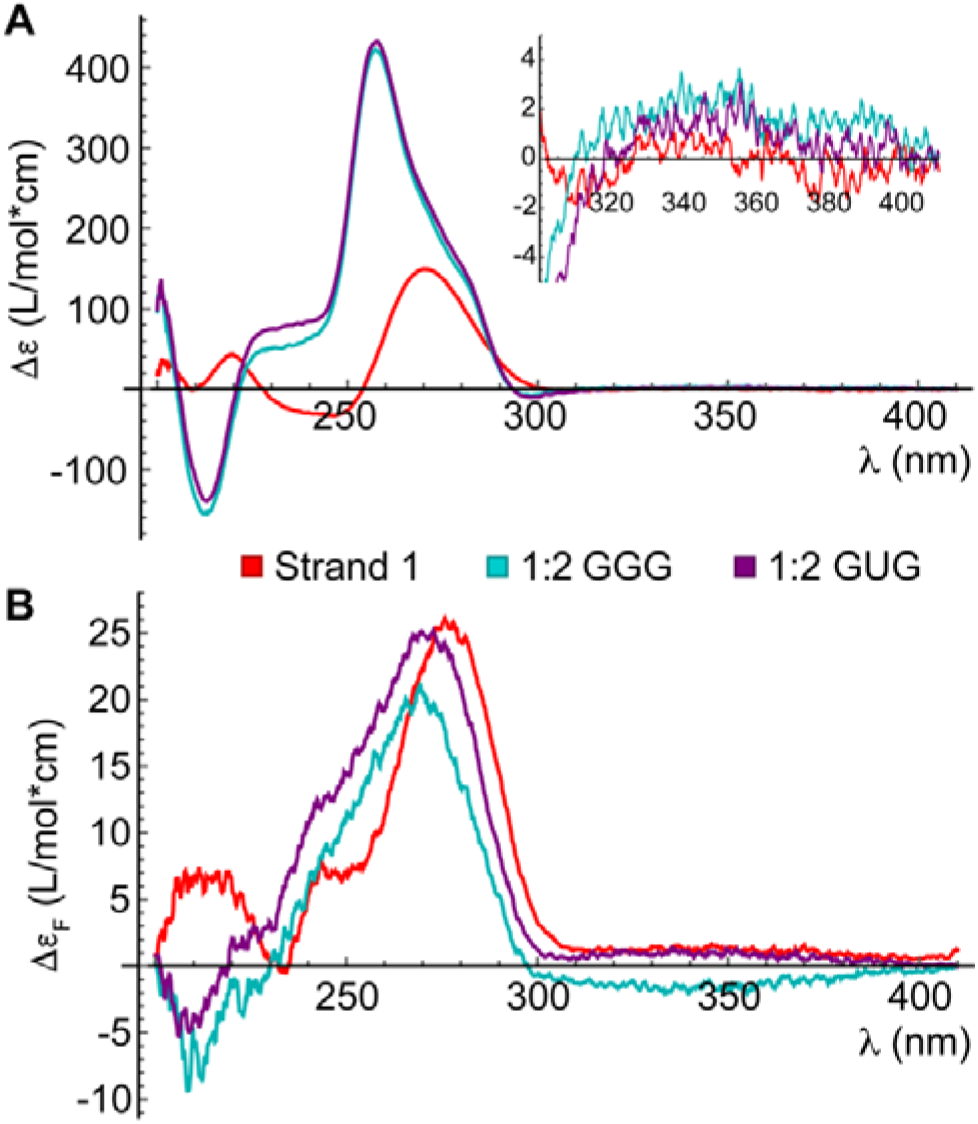
**(A)** CD spectra of ss (red), ds (cyan), and mm (purple) 21-mers. The inset shows a close-up of the 300-410 nm range. **(B)** Processed FDCD spectra of ss (red), ds (cyan), and mm (purple) 21-mers.

We next investigated ss, ds and mm 21-mers at elevated temperature. In all three cases, increasing the temperature leads to a decrease in the overall emission intensity (Fig. S8) and a shortening of the time constant of each decay component (Table 5, Fig. S9). ssRNA shows a population shift from the stacked (longer lifetime) to the unstacked conformation as the temperature is raised. Fully complementary dsRNA shows a population shift from its unique ∼2 ns (presumably base-paired) conformation to a conformation with a lifetime similar to the longer one observed in ssRNA and mm dsRNA. Both duplexes exhibit two transitions in UV melting curves (Fig. S10). A transition at 47 °C for fully complementary dsRNA and 45 °C for mm dsRNA likely corresponds to melting of the AU-rich tails, while a second transition at 65 °C for fully complementary dsRNA and 55 °C for mm dsRNA results in full strand dissociation. Just above its higher T_M_, both duplexes attain nearly identical fluorescence spectra (Fig. S8) and decay properties (Table 5; Figs. 5 and S9) to ssRNA. At all temperatures at which mm dsRNA is intact, its decays are best fit with a single distribution of states (discussed further in Supplementary Note 1). From 30 to 60 °C, the slower decay component of fully complementary dsRNA is best described with a distribution rather than an exponential component, suggesting a disordered ensemble of non-base-paired conformations. The faster decay component of ssRNA is best described with a distribution from 50 to 70 °C (Table 5, Table S3). The fluorescence excitation and emission spectra of mm and fully complementary dsRNA gradually approach those of ssRNA as the temperature is increased (Fig. S8). Specifically, the signature of energy transfer from G disappears from the excitation spectrum and the emission spectrum exhibits a redshift and increase in the red vibronic shoulder characteristic of increased solvent exposure.

## DISCUSSION

In this study, we performed a detailed analysis of the photophysical behavior of pC in RNA using multiple complementary fluorescence-detection methods. Steady-state fluorescence spectroscopy showed that the fluorescence properties of pC in RNA are altered relative to that of monomeric pCr in a sequence-dependent manner. In dinucleotides, the FQY of pC is enhanced relative to pCr when adjacent to an A or C residue and is nearly unchanged when adjacent to a G or U residue. In trinucleotides, the FQY of pC is further enhanced when flanked by A or C residues and enhanced to a much lesser degree when flanked by G or U. The emission spectra of oligonucleotides of all sequences show a consistent hypsochromic shift in emission wavelength and suppression of a long-wavelength vibronic shoulder relative to pCr. The sequence-independent nature of these alterations to the emission spectrum, as well as their appearance when pCr is measured in 30% EtOH, suggest that they predominantly reflect the polarity of the local environment experienced by pC. Indeed, the extent of the changes in both λ_max_ and lineshape in fully aqueous solvent is correlated to the degree of protection from the solvent, where the spectra of dinucleotides deviate the least from pCr and the spectrum of fully complementary dsRNA deviates the most (dsRNA ≈ mismatched duplex >> ssRNA ≈ CpCC ≈ pCC >> pCr). In contrast, 2-AP experiences minimal changes in lineshape or λ_max_ across the analogous series of 2-APr, (2-AP)C and C(2-AP)C (Fig. S11) (11). Observation of inter-strand energy transfer in fluorescence excitation spectra of pC further differentiates double-stranded from single-stranded environments. Facile measurement of steady-state fluorescence spectra can therefore give a general identification of the local environment of pC in RNA of unknown structure.

While comparison to 2-AP is illuminating, other FBAs exhibit properties more similar to pC such as moderate variation in FQY and environment-sensitive emission spectra. A standout among “isomorphic” FBAs (those that at least roughly retain the size and shape of their cognate native bases) is ^th^G (39). It has been widely used as a probe of conformational changes (40, 41), including those induced by protein binding (42–45) and catalysis (46), and as a scaffold for the development of more specialized FBA-based probes (47, 48). The emission of ^th^G is sensitive to both base stacking, which induces a moderate reduction in FQY from 0.46 for the free nucleoside to ∼0.1 when flanked by Gs or Ts, and base pairing (39, 49).This ∼5-fold difference in FQY is similar to the contrast we observe between the brightest and darkest pC-labeled oligonucleotides, though the maximum FQY we observe for pC (0.09) is substantially lower.

The most widely used and studied fluorescent cytosine analogues include an expansive family of tricyclic probes such as tC (50), tC^0^ (51), DEA-tC (52), and ABN (53). tC and tC^0^ are noteworthy for having nearly identical FQY (∼0.2) as free nucleosides and in single-stranded and double-stranded nucleic acids (50, 51), while DEA-tC has the unusual property of being brightest in double-stranded nucleic acids (52, 54). The minimally fluorescent tC_nitro_ (55), has been used as a FRET acceptor with tC^0^ as the donor (56), and the exceptionally bright ABN is readily detectable at the single molecule level (53). pC exists in a space complementary to these advanced FBAs, with its small size and environment-sensitive fluorescence justifying its selection when a general probe of nucleic acid structure and/or protein interactions is desired.

Time-resolved fluorescence measurements demonstrated that pC typically samples two distinct environments with different emissive properties in short RNA oligonucleotides. This stands in contrast to previous measurements in which the decays of the DNA dinucleotides deoxyribo-pCT (dpCT), dpCC, dpCA, dpCG and trinucleotide dTpCT were all adequately fit with mono-exponential models, and only dGpCG required a bi-exponential model (17). The sequence dependence of both intensity and lifetime observed in DNA was far more mild than we observe in RNA (17). The previously studied DNA oligonucleotides may interconvert between stacked and unstacked conformations at a rate on the order of or faster than the relaxation of pC, which would prevent their resolution into distinct decay components. Both computational (57) and experimental (58) results have suggested that this factor impacts the fluorescence decays of 2-AP in short DNA oligonucleotides. It is also possible that, in contrast to our observations in RNA, pC has similar lifetimes in the stacked and unstacked conformations of certain DNA dinucleotides. Indeed, it was proposed based on molecular dynamics simulations that dpCT exists on average in a stacked conformation, suggesting that a T residue stacked on only one side of pC is insufficient to alter its photophysics (17). Our observation that the decays of pCU and UpCU are best fit with a tri-exponential or distribution model indicates that they lack a single, well-defined base-stacked structure. This may reflect the fact that base-stacked structures of U-containing oligonucleotides are generally less stable than those of other native bases (59–61).

Through comparison of the fluorescence decays of different RNA species, comparison of fully aqueous solvent to 30% ethanol, and comparison of CD and FDCD spectra, we provide strong evidence that the brighter, more slowly decaying species occupies a base-stacked structure. FDCD previously revealed the exact opposite for 2-AP, with the bright conformations being so drastically unstacked that their CD spectra resemble free 2-AP riboside (11). The bright conformations that FDCD reports on additionally exhibit, with the exception of pCU, efficient energy transfer from native bases to pC. We note that a previous publication reported attempting FDCD measurements on pC-labeled DNA (in a TpCA sequence context) in the 300-420 nm range, but no signal was observed with the instrumentation used in that study even at higher concentrations than those used here (62).

The enhancement of pC fluorescence in trinucleotides of all sequence contexts compared to the corresponding dinucleotides is accompanied by both a shift toward the stacked conformation and a slowing of this conformation’s decay rate, suggesting that the stacked conformation is both more prevalent and more emissive in trinucleotides. Previous computational studies have found that the S_0_ □ S_1_ transition of pC exhibits reduced oscillator strength when it is base-stacked with G on one or both sides in the geometry of an A- or B-form helix, reducing the predicted rate of radiative relaxation (10, 18, 63). Our computational studies replicate this finding and show that it holds to a slightly lesser degree in CpCC, which is consistent with the experimentally observed decrease in the molar absorptivity of pC in trinucleotides relative to pCr (Table 2). This suggests that the enhancement of pC fluorescence observed in both DNA (17) and RNA trinucleotides relative to a pC monomer originates from a decrease in the overall rate of nonradiative decay. The rigidity and solvent protection afforded to pC in a trinucleotide as compared to pCr, combined with the conspicuous absence of low-lying charge transfer or dark states, could enable the non-radiative rate to decrease in trinucleotides to a greater degree than the radiative rate, giving rise to the observed enhancement of fluorescence.

It is worth noting that among pC monomer-containing oligonucleotides, sub-nanosecond components were found only for the 3E and 1E+1G fits to the decays of pCU and UpCU in buffer, respectively (Tables 2 and S2). This suggests that our benchtop TCSPC instrumentation is sufficient to resolve all decay components of the pC-containing oligonucleotides investigated. If the decays of all conformations are indeed being successfully resolved, the intensity ratio between pCr and an oligonucleotide should be roughly equal to the ratio of their ⟨τ⟩_obs_ values (64). In Table 6, these ratios are compared for two pC- and 2-AP-labeled oligonucleotides of analogous sequence, measured under identical experimental conditions. It is clearly evident that the observed changes in lifetime are adequate to explain the changes in intensity of these pC-containing oligonucleotides, while the same is not true of 2-AP (11). This is consistent with the repeated observation of decay components in the 10s of picoseconds or shorter for 2-AP-containing oligonucleotides in ultrafast measurements (65, 66). Thus, our results suggest that pC can be more thoroughly characterized with readily accessible instrumentation than 2-AP, a benefit that was also noted when oligonucleotides containing ^th^G were compared to equivalent sequences containing 2-AP (67).

**Table 6.** FQY and observed average lifetime ratios (oligonucleotide : riboside) for 2- AP and pC.

|  | **X=2-AP<sup>1</sup>** |  | **X=pC<sup>2</sup>** |  |
| --- | --- | --- | --- | --- |
| **Sequence** | **FQY/FQY<sub>riboside</sub>** | **$\langle\tau\rangle_{\text{obs}}/\langle\tau\rangle_{\text{obs,riboside}}$** | **FQY/FQY<sub>riboside</sub>** | **$\langle\tau\rangle_{\text{obs}}/\langle\tau\rangle_{\text{obs,riboside}}$** |
| XC | 0.25 | 0.55 | 2.35 | 2.25 |
| CXC | 0.10 | 0.44 | 3.49 | 3.56 |

<sup>1</sup>Recalculated from data in ref. 11
<sup>2</sup>This work

pC proved to be a sensitive probe of base-pairing, as previously observed in both systematic investigation of model DNA oligonucleotides (17) and benchmarking using variants of biologically relevant RNA (4). We found that a fully complementary duplex exhibited significantly reduced emission intensity relative to ssRNA and mismatched dsRNA, showing that base-pairing facilitates nonradiative relaxation of pC. Tautomerization in the excited state via double proton transfer between pC and G has been identified computationally as a plausible quenching mechanism (68). We further observed a dominant ∼2 ns decay component unique to fully complementary dsRNA, which is comparable to the reported average lifetimes of pC in a variety of dsRNA fragments designed to mimic segments of a T-box riboswitch-tRNA complex (4). We observed that dsRNA and mismatched duplex have very similar emission spectra that indicate that pC occupies a similarly nonpolar environment in both cases, while ssRNA and mismatched duplex have similar emission intensities and decays (Fig. 5). The quenching of pC in fully complementary dsRNA therefore seems to be a direct consequence of base-pairing rather than the protected and base-stacked structural context of dsRNA.

In this work, we take a multi-pronged approach to empirically characterize the structure-sequence-photophysics relationships of pC in RNA. Further experimental and computational investigation is required to identify the precise photophysical mechanisms underlying the results reported here. Previous theoretical studies of base-stacked complexes have identified mechanisms by which pC could be quenched by G, such as electron transfer from pC (17, 18) and reductions in the oscillator strength of the fluorescence transition (10, 63), but do not explain the overall enhancement of pC fluorescence observed in GpCG. While our computational results also do not reveal an obvious source of fluorescence enhancement, even in CpCC, they do show that certain quenching mechanisms that impact other FBAs are absent in pC. In future work, we plan to dissect the relaxation pathways that occur prior to emission by utilizing ultrafast fluorescence upconversion spectroscopy to measure time-resolved emission spectra at a sub-picosecond timescale.

The wide-ranging empirical characterization reported here will facilitate informed design of biologically relevant pC-containing RNA sequences and nuanced interpretation of the ensuing findings. We demonstrate that steady-state emission spectra reflect the polarity of the local environment of pC, while the energy transfer observed in excitation spectra distinguishes single-from double-stranded environments. TCSPC resolves heterogeneity in base stacking and hydrogen bonding and FDCD confirms that base-stacked structures are generally brighter and exhibit efficient energy transfer. The findings reported here will be applied to further develop pC as a reporter of RNA conformational changes in response to environmental stimuli.

## DATA AVAILABILITY

The data underlying this article will be shared on reasonable request to the corresponding author.

## SUPPLEMENTARY DATA

Supplementary Data are available at NAR online.

## AUTHOR CONTRIBUTIONS

T.L.C.: Conceptualization, Investigation, Visualization, Formal Analysis, Writing - original draft. J.R.W.: Conceptualization, Investigation, Visualization, Supervision, Funding Acquisition, Writing - review and editing.

## ACKNOWLEDGEMENTS

The authors acknowledge Dr. Jim Prell of the University of Oregon and lab member Austin Green for contributing mass spectrometry instrumentation and expertise.

