## Supplemental information for "Impacts of sequence and structure on pyrrolocytosine fluorescence in RNA"

Includes:

Supplementary Figures S1-S11

Supplementary Note 1: Comparison of fitting models for TCSPC data

Supplementary Note 2: Interpretation of $g_F$ spectra obtained through FDCD

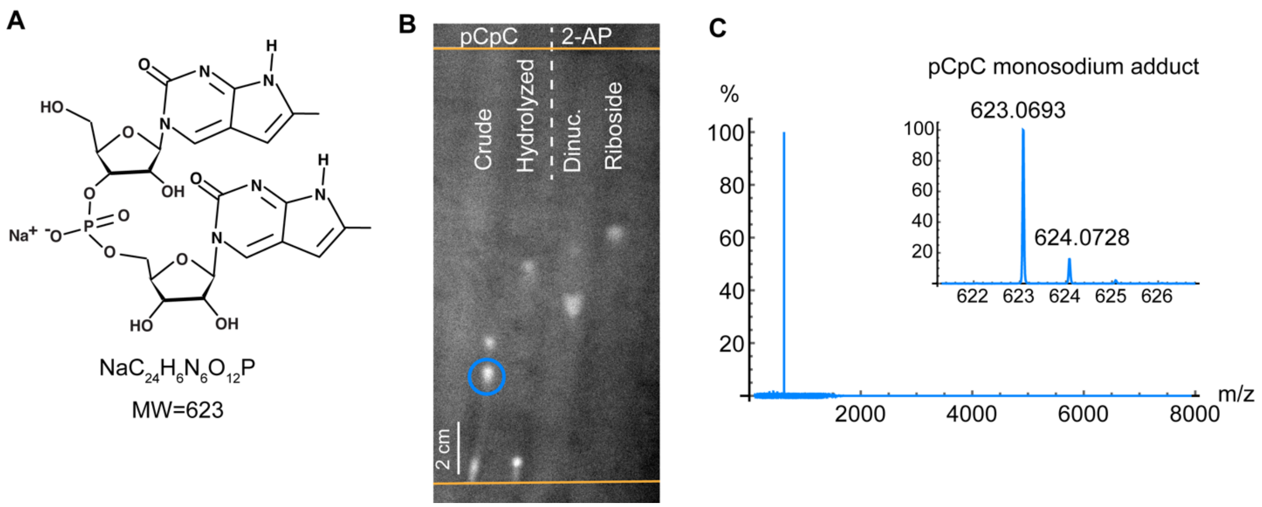


**Figure S1.** Preparation of pC dinucleotide. (**A**) Structure and chemical formula of pCpC. (**B**) Analytical TLC plate imaged on an ultraviolet transilluminator after two elutions (details in methods section). The origin and solvent front are indicated by orange lines. Two spots disappear when pCpC is base-hydrolyzed, suggesting that they contain sugar-phosphate linkages. Following purification by preparative TLC with the same elution procedure, the lower spot (blue circle) was identified by mass spectrometry as pure pCpC. 2-AP dinucleotide and riboside are included as references, demonstrating that dinucleotides migrate more slowly than the corresponding nucleosides. (**C**) Mass spectrum of pCpC purified by preparative TLC. Inset: Zoom-in of the region that shows signal.

| pCpC | | 2-AP | |
|---|---|---|---|
| Crude | Hydrolyzed | Dinucleotide | Nucleoside |
| 0.03 | 0.03 | 0.37 | 0.48 |
| 0.18 | 0.31 | 0.46 | -- |
| 0.27 | 0.42 | -- | -- |

**Table S1.** Retention factor values of all resolvable spots in Fig. S1b.

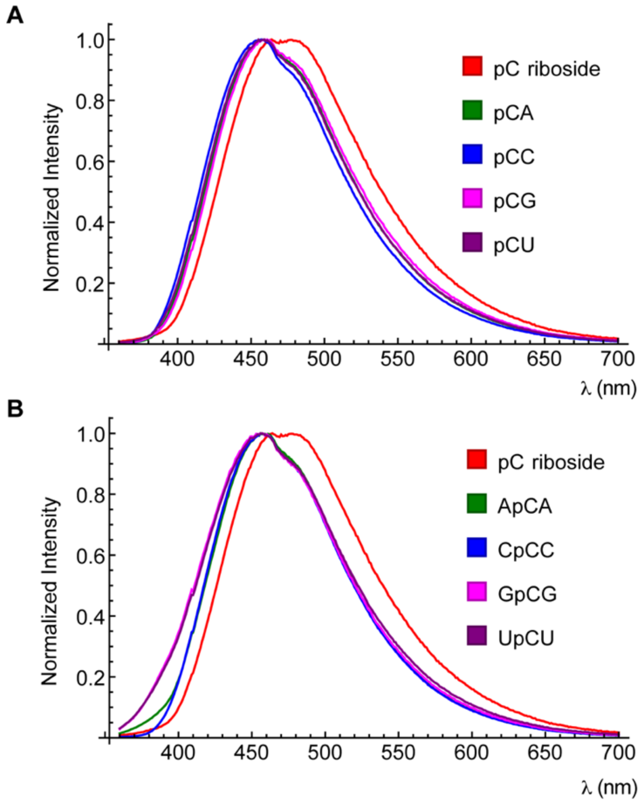


**Figure S2. (A)** Emission spectra of pCr and all dinucleotides normalized to a peak value of 1. **(B)** Emission spectra of pCr and all trinucleotides normalized to a peak value of 1. A blue-shift in peak emission wavelength and suppression of a long-wavelength shoulder is observed in all sequence contexts.

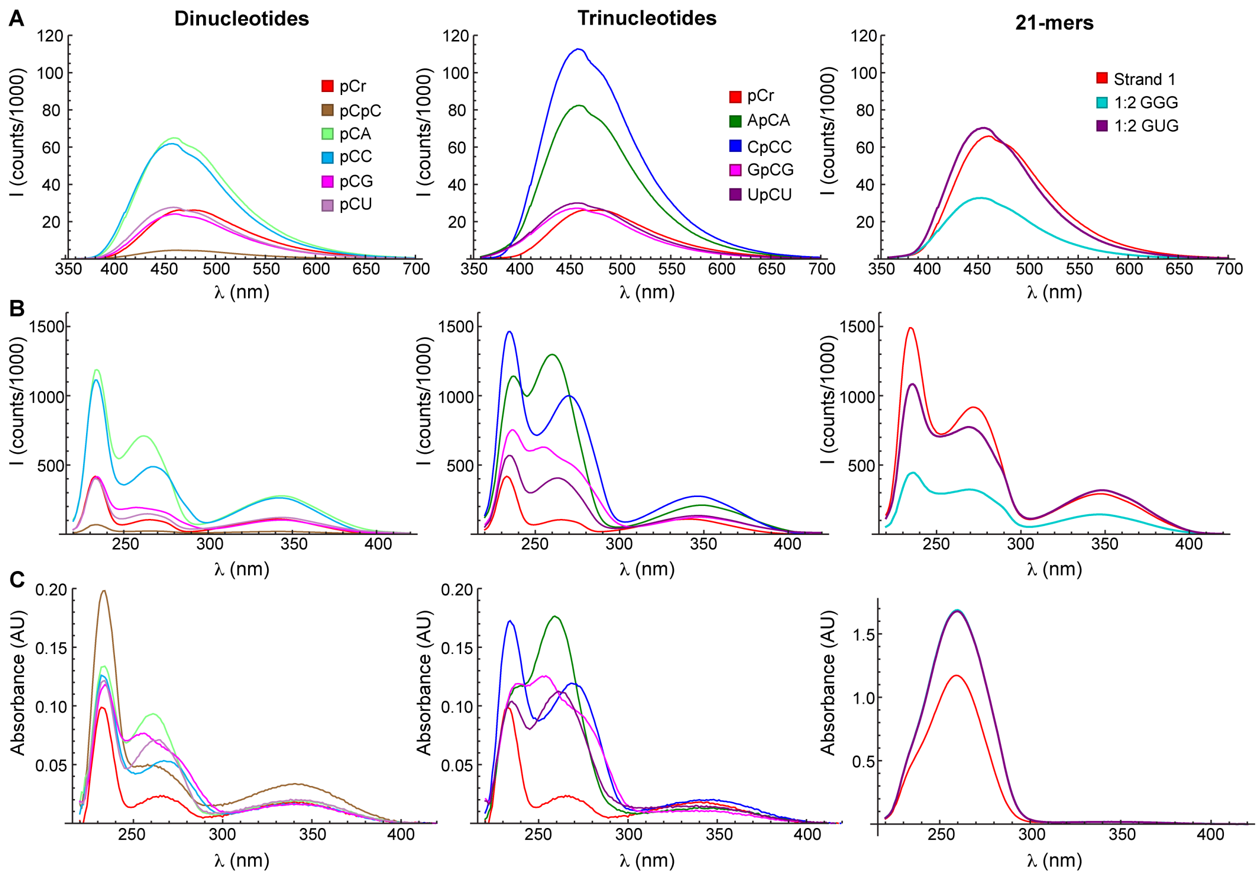


**Figure S3.** Un-normalized steady-state spectra of dinucleotides (left column), trinucleotides (middle column) and 21-mers at 20 °C. (**A**) Fluorescence emission spectra under excitation at 340 nm. (**B**) Fluorescence excitation spectra recorded with detection at 460 nm. (**C**) Absorbance spectra.

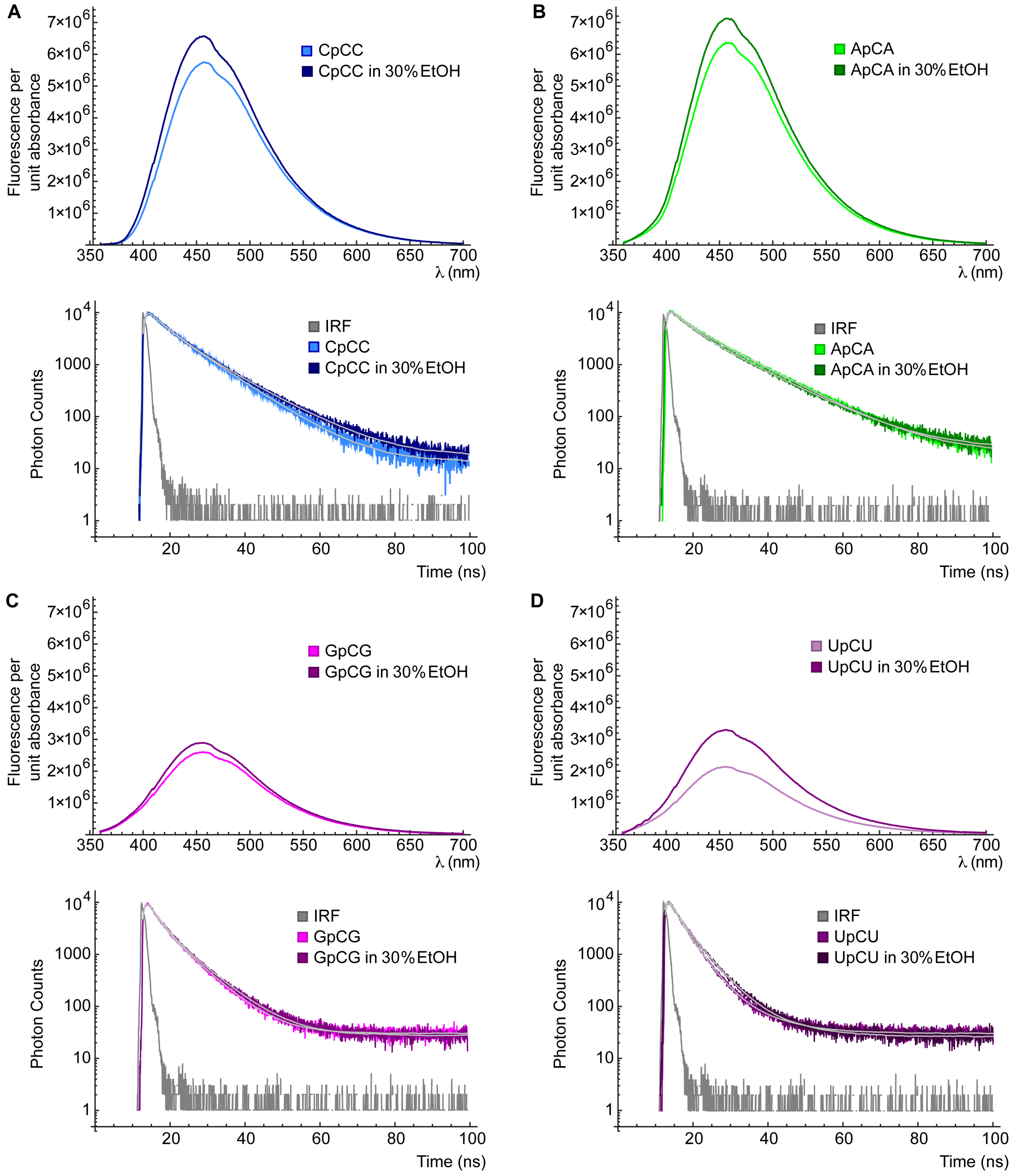


**Figure S4.** Comparison of fluorescence emission spectra and TCSPC of trinucleotides in buffer (lighter colors) and buffer containing 30% ethanol (darker colors). Multi-exponential fits (tri-exponential for UpCU in buffer, bi-exponential for all others) are overlaid in gray. (**A**) CpCC, reproduced from main text Figure 3. (**B**) ApCA. (**C**) GpCG. (**D**) UpCU.

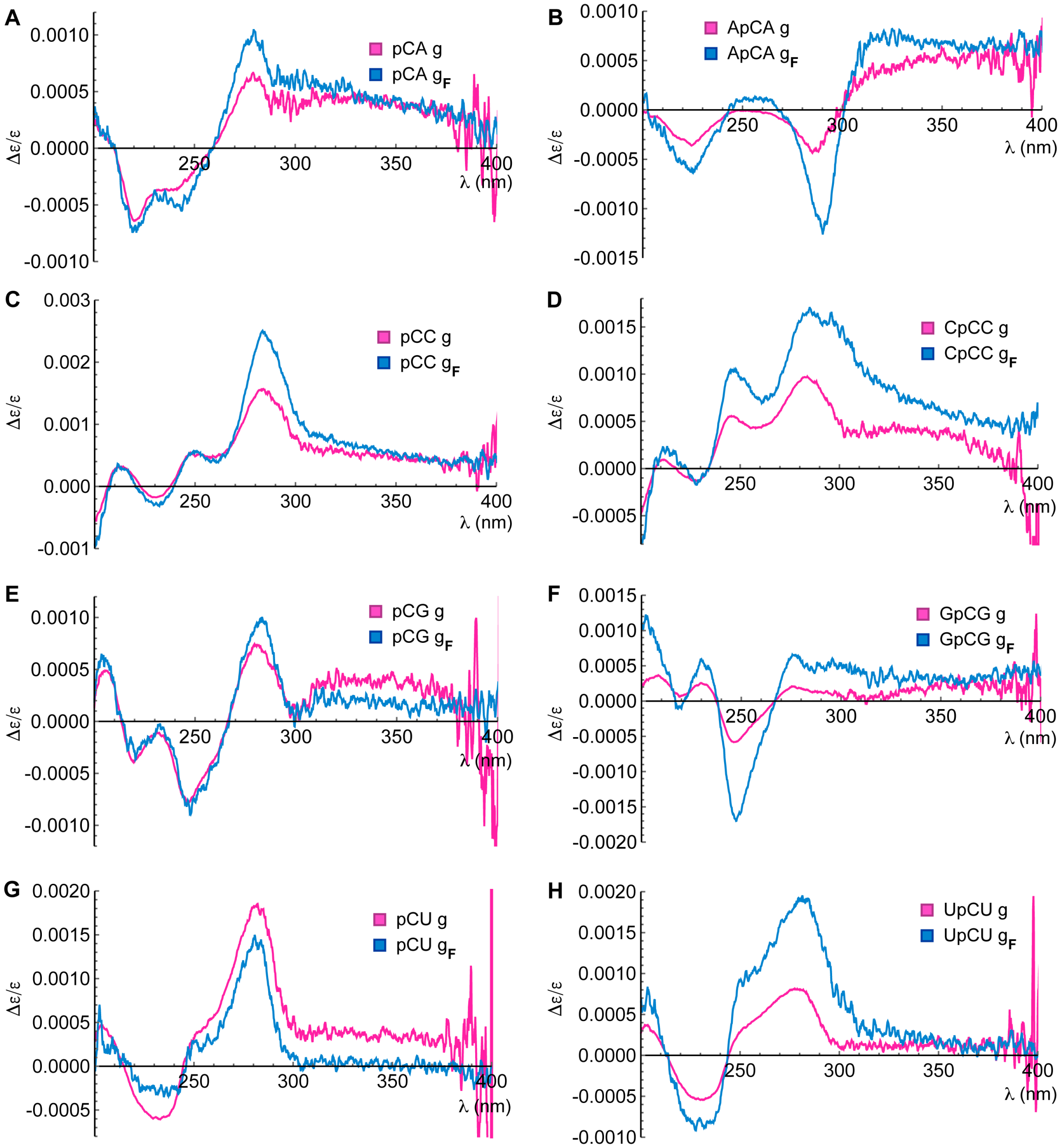


**Figure S5.** g (pink) and $g_F$ (blue) spectra of di- and trinucleotides. **(A)** Spectra of pCA. **(B)** ApCA. **(C)** pCC. **(D)** CpCC. **(E)** pCG. **(F)** GpCG. **(G)** pCU. **(H)** UpCU.

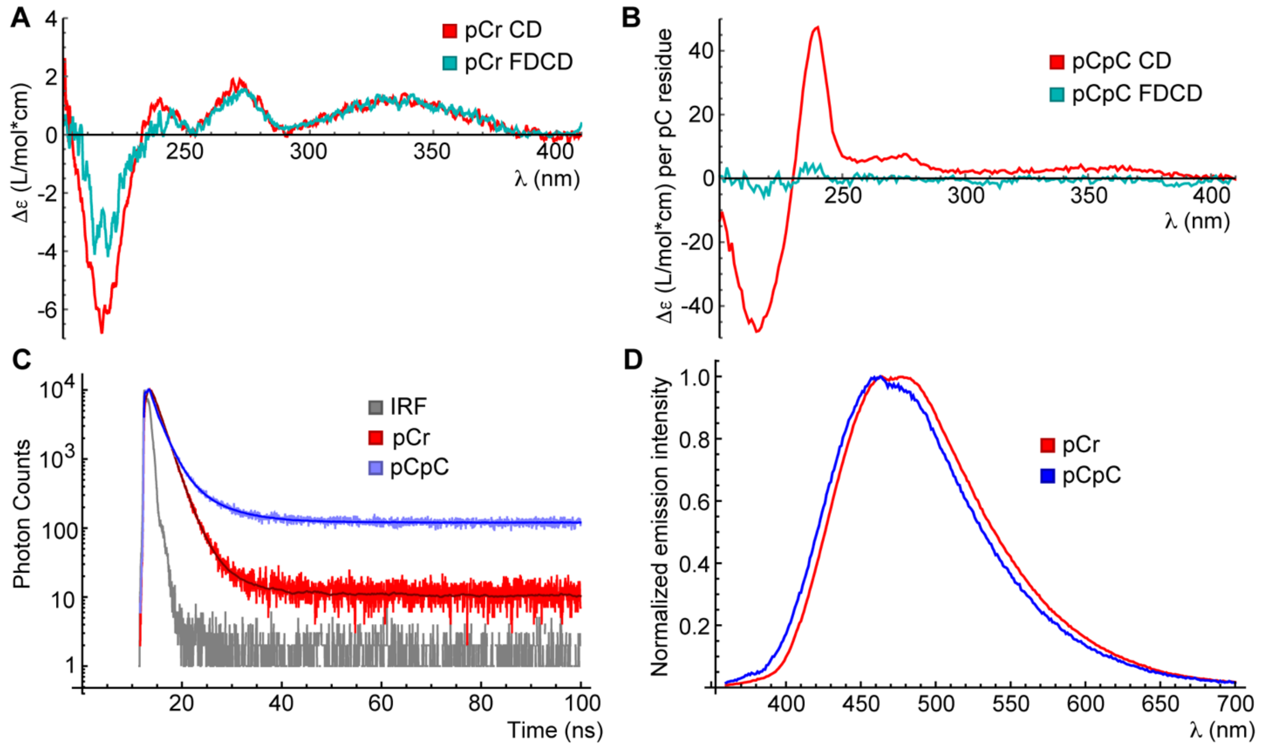


**Figure S6.** Spectroscopy of pCr and pCpC. **(A)** CD (red) and processed FDCD (cyan) spectra of pCr. **(B)** CD (red) and processed FDCD (cyan) spectra of pCpC. **(C)** TCSPC of pCr (red), pCpC (blue), and instrument response function (gray). Fits are overlaid in a darker shade. **(D)** Normalized emission spectra of pCr (red) and pCpC (blue).

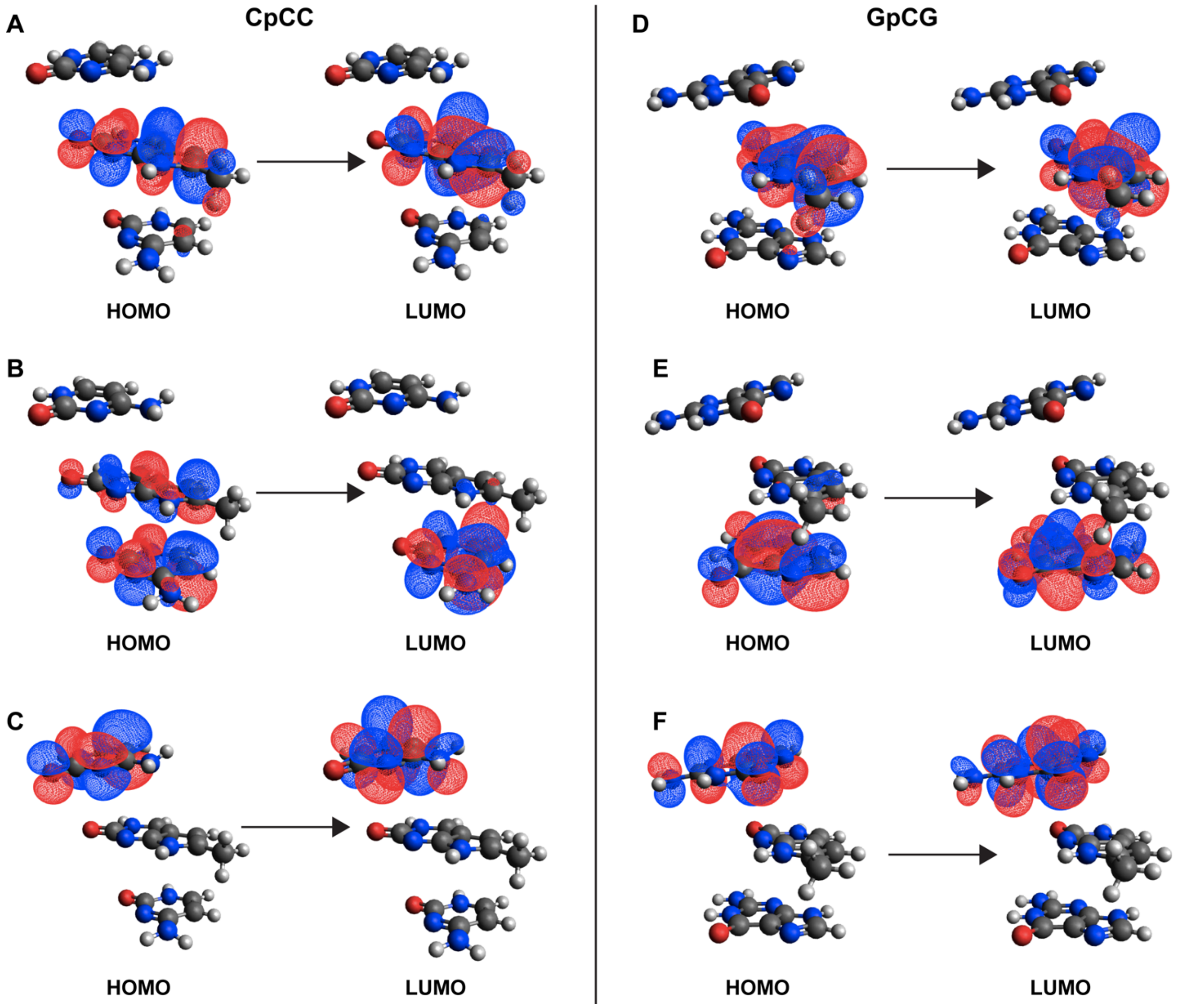


**Figure S7.** Natural transition orbitals for excited states of interest in CpCC (left column) and GpCG (right column). (**A**) CpCC $S_1$ (**B**) CpCC $S_4$ (**C**) CpCC $S_5$ (**D**) GpCG $S_1$ (**E**) GpCG $S_8$ (**F**) GpCG $S_9$

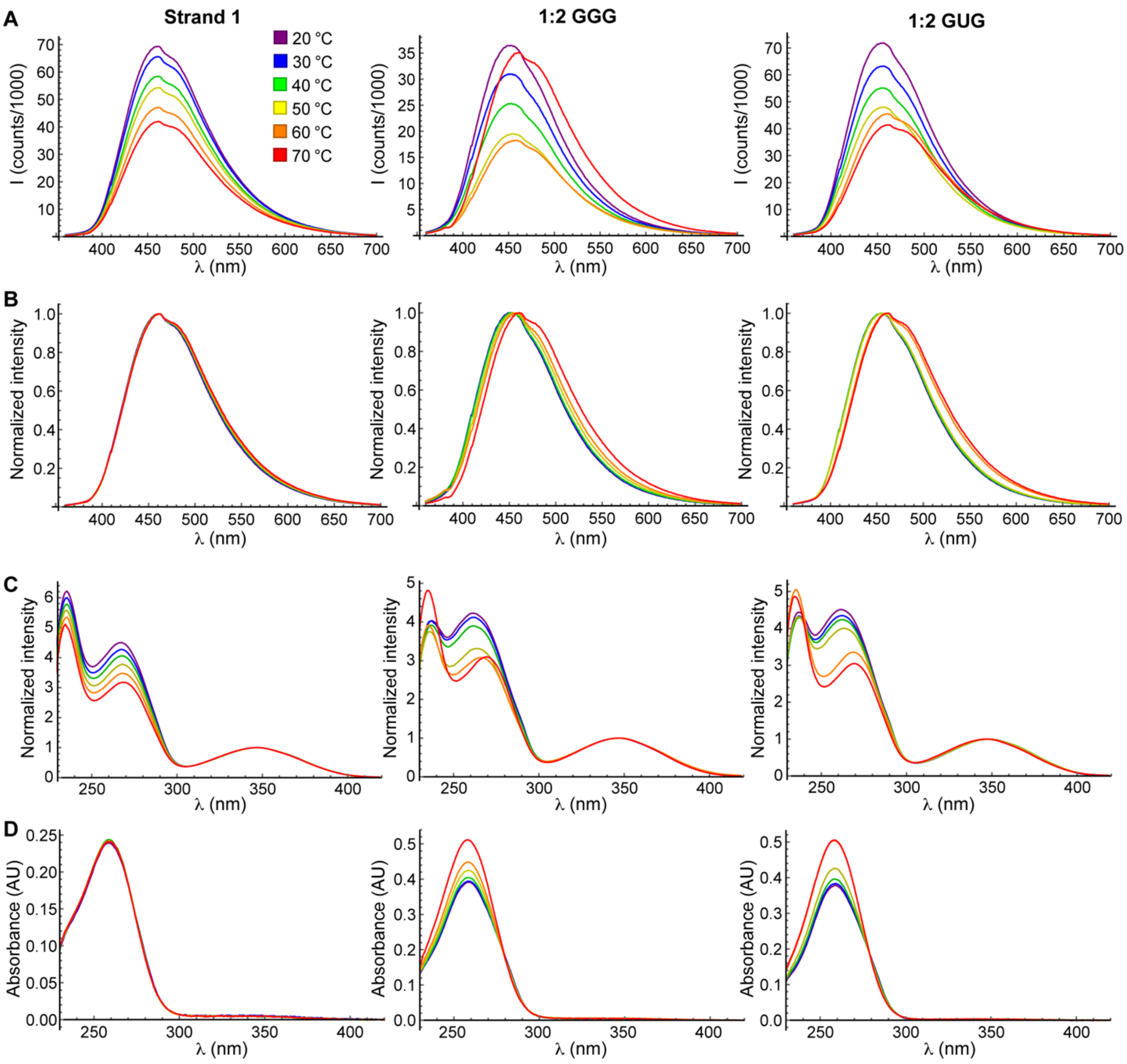


**Figure S8.** Steady-state spectra of ss (left column) ds (middle column) and mm (right column) 21-mers at 20 °C (purple), 30 °C (blue), 40 °C (green), 50 °C (yellow), 60 °C (orange) and 70 °C (red). **(A)** Fluorescence emission spectra under excitation at 340 nm. **(B)** Spectra from row A normalized to a peak intensity of 1. **(C)** Fluorescence excitation spectra recorded with detection at 460 nm with the longest-wavelength peak normalized to an intensity of 1. **(D)** Absorbance spectra.

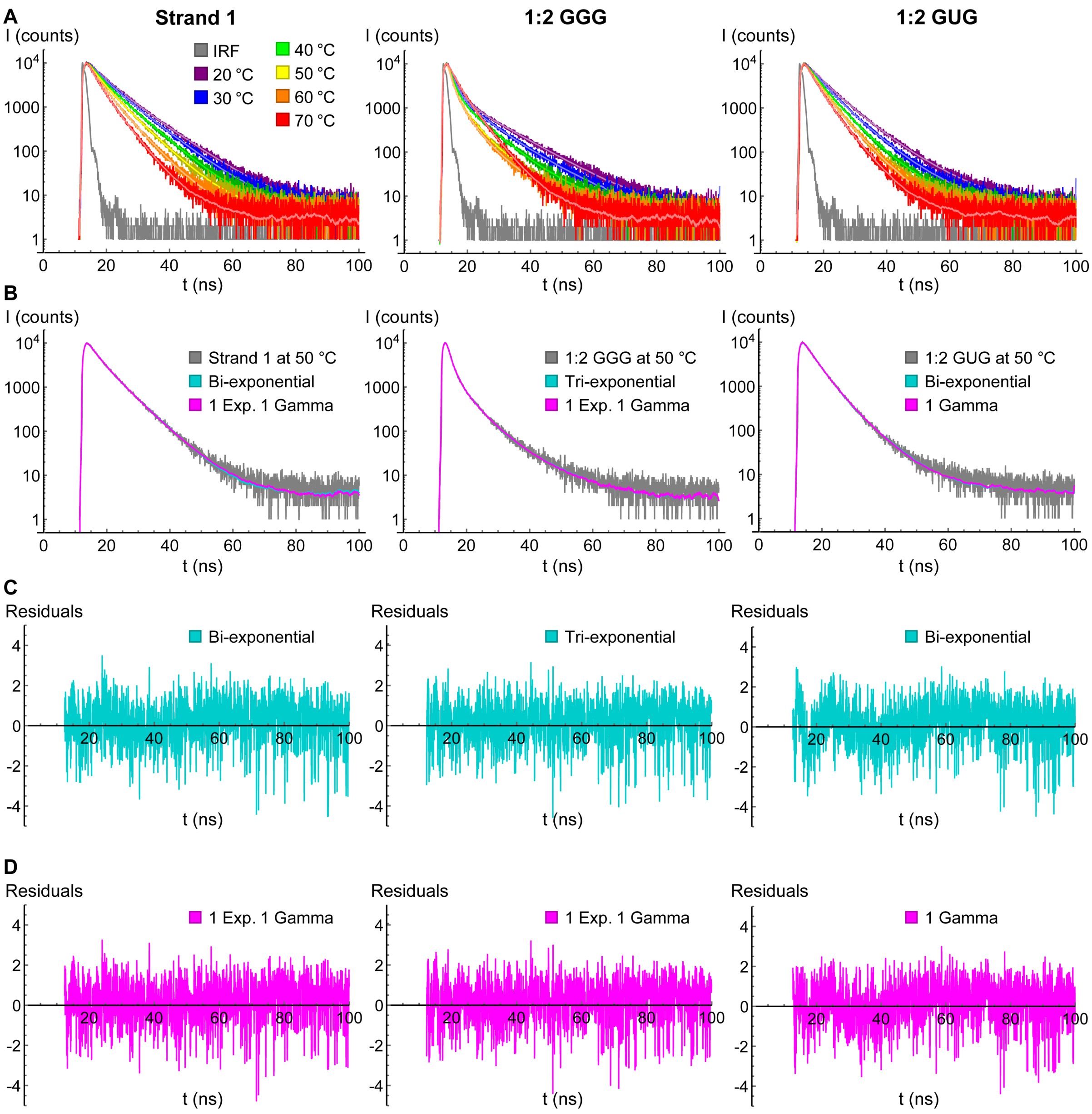


**Figure S9.** TCSPC of ss (left column) ds (middle column) and mm (right column) 21-mers. **(A)** Fluorescence decays recorded at 20 °C (purple), 30 °C (blue), 40 °C (green), 50 °C (yellow), 60 °C (orange) and 70 °C (red). Fits (details in Table 5) are overlaid in lighter color. A typical IRF is plotted in gray. (B-D) Comparison of multiexponential and distribution fits. Data recorded at 50 °C are used as an example. **(B)** Decays and fits. Left: Biexponential (cyan, $\chi^2$=1.21) and 1 exponential + 1 gamma (magenta, $\chi^2$=1.16) fits to strand 1 decay (gray). Middle: Triexponential (cyan, $\chi^2$=1.19) and 1 exponential + 1 gamma (magenta, $\chi^2$=1.18) fits to 1:2 GGG decay (gray). Right: Biexponential (cyan, $\chi^2$=1.29) and 1 gamma (magenta, $\chi^2$=1.22) fits to 1:2 GUG decay (gray). **(C)** Residuals from the multiexponential fits shown in row B. **(D)** Residuals from the distribution fits shown in row B. The residuals for the 1 gamma fit of 1:2 GUG show some structure, but triexponential ($\chi^2$=1.16) and 1 exponential + 1 gamma ($\chi^2$=1.20) fits showed limited improvement over 1 gamma ($\chi^2$=1.22).

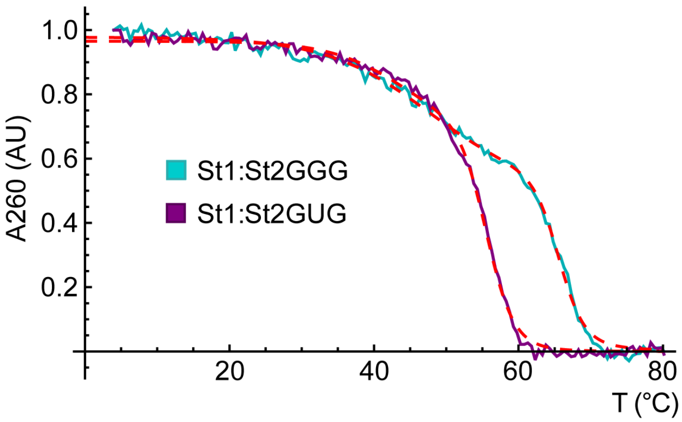


**Figure S10.** Melting curves of ds (cyan) and mismatched (purple) 21-mers fit with a two-transition model (dashed red). The ds 21-mer shows transitions at 47.1 and 65.4 °C, and the mismatched 21-mer shows transitions at 44.5 and 55.5 °C.

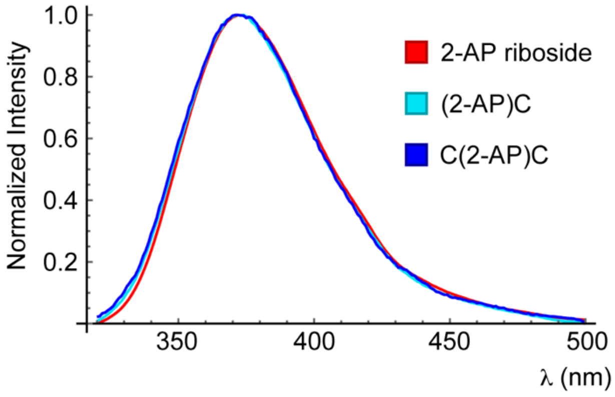


**Figure S11.** Normalized fluorescence emission spectra of 2-aminopurine riboside (red), (2-AP)C (cyan), and C(2-AP)C (blue). Data re-processed from ref. 11.

**Supplementary Note 1: Comparison of fitting models for TCSPC data**

TCSPC data for most oligonucleotides and measurement conditions were fit very well ($\chi^2 < 1.2$) with bi-exponential (2E) models. However, in certain cases 2E fits were poor or only moderately good:

<u>Moderately good ($1.2 < \chi^2 < 1.3$)</u>
pCU
UpCU in buffer
St1 at 50, 60 and 70 °C
St1:St2GGG at 30 and 70 °C
St1:St2GUG at 30, 40, 50 and 70 °C

<u>Poor ($\chi^2 > 1.3$)</u>
pCpC
St1:St2GGG at 40, 50 and 60 °C

Tri-exponential (3E) fits offered significant improvement in $\chi^2$ over 2E for the scenarios listed above as poor and offered modest improvement for those listed as moderately good. In some cases, however, 3E fits led to parameter values that were weakly constrained in the fit, and/or had poor reproducibility across replicates. For example, a 3E fit to the decay of St1:St2GGG at 30 °C introduced a component with time constants of 11,12 and 15 and weights of 7, 4 and 2% across three replicates. As a result, we investigated whether a previously reported distribution model (details in methods section) containing either a single gamma component (1G) or one gamma and one exponential component (1E+1G) could better describe this data. A 1G model requires 2 parameters to describe the molecular properties underlying the observed decay, 2E requires 3, 1E+1G requires 4, and 3E requires 5. Each gamma component contributes 3 free parameters (w, $\alpha$ and $\kappa$) and each exponential component contributes 2 (w and $\tau$), with the constraint that the weights add up to 1 reducing the number of parameters by 1.

Table S2 compares 2E, 1E+1G and 3E (reproduced from Table 1) models for pCU and UpCU. For pCU, the 4-parameter 1E+1G fit closely mimics the 5-parameter 3E fit, but with the two longer components of the 3E fit merged into one distribution. The fact that these two components can be merged into one distribution without $\chi^2$ suffering suggests that there might not be three distinct stable states that persist for long enough to resolve into separate components. The parallels between the 1E+1G and 3E fits for UpCU are less quantitative, but in this case too the gamma component has a time constant between the two slower components of the 3E fit and a weight larger than either of the 3E ones alone. Together, these fits indicate that pCU and UpCU sample a more complex ensemble of structures than the two that adequately describe other di- and trinucleotides, and/or that adjacent U residues make the photophysics of pC more complex.

Interestingly, a 2E model describes the decay of UpCU in buffer containing 30% EtOH very well ($\chi^2$=1.14). When either of the high-quality fits to its decay in fully aqueous buffer (1E+1G or 3E) are compared to its high-quality 2E fit in EtOH, UpCU displays the shift in population to the shorter component that was expected to accompany unstacking. However, this shift is not observed if fits from the same model are compared across buffer and 30% EtOH, highlighting the complexity that adjacent U residues confer onto pC's structure-photophysics relationship.

| RNA | **Model** | $\chi^2$ | **$\langle\tau\rangle_{obs}$ (ns)** | **$w_1$** | **$\tau_1$ (ns)** | **$w_2$** | **$\tau_2$ or $\langle\tau_2\rangle$ (ns)** | **$w_3$** | **$\tau_3$ (ns)** |
|---|---|---|---|---|---|---|---|---|---|
| pCU | 2E | 1.25 | 2.25 | 0.80 | 1.56 | 0.20 | 5.01 | | |
| pCU | 1E+1G | 1.08 | 2.06 | 0.56 | 1.04 | *0.43 | *3.36 | | |
| pCU | 3E | 1.08 | 2.01 | 0.54 | 0.98 | 0.37 | 2.8 | 0.09 | 6.3 |
| UpCU | 2E | 1.22 | 3.46 | 0.8 | 2.76 | 0.2 | 6.2 | | |
| UpCU | 1E+1G | 1.09 | 2.99 | 0.16 | 0.54 | *0.83 | *3.48 | | |
| UpCU | 3E | 1.06 | 3.19 | 0.33 | 1.4 | 0.64 | 3.4 | 0.04 | 9.5 |
| [1]UpCU | 2E | 1.14 | 4.06 | 0.68 | 3.00 | 0.32 | 6.28 | | |
| [1]UpCU | 1E+1G | 1.07 | 3.84 | 0.14 | 1.15 | *0.86 | *4.28 | | |
| [1]UpCU | 3E | 1.06 | 3.91 | 0.33 | 1.88 | 0.63 | 4.6 | 0.05 | 10 |

Table S2. Comparison of 2E, 1E+1G and 3E fits for pCU and UpCU. Mean values across three successive data acquisitions.
*Parameter originates from a gamma component
[1]Data recorded in buffer containing 30% EtOH

Distribution models were also investigated for the ss, ds and mm 21-mers. For these samples, we found that 1G or 1E+1G models were better justified (lower $\chi^2$ or comparable $\chi^2$ but fewer parameters) than 3E for all the conditions in which 2E fits were moderately good or poor. For example, Table S3 presents the results of 2E, 1E+1G and 3E fits to the decays of St1 at temperatures of 50, 60 and 70 °C, where 2E fits were only moderately good ($1.2 < \chi^2 < 1.3$). 1E+1G fits offer a clear improvement over 2E fits, while 3E fits exhibit slightly poorer $\chi^2$ than 1E+1G despite having more free parameters. Analogous results were observed for the fully complementary St1:St2GGG at temperatures of 30, 40, 50, 60 and 70 °C, with $\chi^2$ being marginal or poor with a 2E model and slightly better with a 1E+1G model than a 3E model. This led us to conclude that a 1E+1G model was best justified for those conditions.

| RNA | °C | Model | $\chi^2$ | $\langle\tau\rangle_{obs}$ (ns) | $w_1$ | $\tau_1$ or $\langle\tau_1\rangle$ (ns) | $w_2$ | $\tau_2$ (ns) | $w_3$ | $\tau_3$ (ns) |
|---|---|---|---|---|---|---|---|---|---|---|
| St1 | 50 | 2E | 1.21 | 4.70 | 0.52 | 2.97 | 0.48 | 6.61 | | |
| | 50 | 1E+1G | 1.16 | 4.62 | 0.61* | 3.54* | 0.39 | 6.32 | | |
| | 50 | 3E | 1.17 | 4.66 | 0.44 | 2.61 | 0.56 | 6.15 | 0.01 | 30 |
| | 60 | 2E | 1.25 | 3.97 | 0.56 | 2.58 | 0.44 | 5.74 | | |
| | 60 | 1E+1G | 1.16 | 3.89 | 0.65* | 3.06* | 0.35 | 5.48 | | |
| | 60 | 3E | 1.19 | 3.90 | 0.40 | 2.06 | 0.55 | 4.91 | 0.04 | 8.80 |
| | 70 | 2E | 1.23 | 3.42 | 0.62 | 2.42 | 0.38 | 5.07 | | |
| | 70 | 1E+1G | 1.17 | 3.38 | 0.73* | 2.78* | 0.27 | 4.87 | | |
| | 70 | 3E | 1.20 | 3.35 | 0.53 | 2.18 | 0.46 | 4.69 | 0.004 | 12.72 |

Table S3. Comparison of 2E, 1E+1G and 3E fits for the ss 21-mer. Mean values across three successive data acquisitions.
*Parameter originates from a gamma component

The mismatched 21-mer, St1:St2GUG, exhibits slightly different behavior. While its decay was well fit with a 2E model at 20 °C ($\chi^2$=1.13), the fit was slightly better with a 1G model ($\chi^2$=1.11) despite the latter having fewer free parameters. This stands in contrast St1 at 20 °C, which is fit considerably better with a 2E model ($\chi^2$=1.12) than 1G ($\chi^2$=1.23). This distinction between ss and mm RNAs becomes even more prominent at elevated temperature, with a 1G model yielding $\chi^2$=1.49 for ssRNA and 1.22 for mm dsRNA at 30 °C. Furthermore, at 50 °C, just below its melting temperature, the mm 21-mer's decay is better fit by a 1G model ($\chi^2$=1.22) than a 2E model ($\chi^2$=1.29). At 60 °C, just above melting, it joins St1 in being considerably better fit by a 2E or 1E+1G model ($\chi^2$=1.14 and 1.09, respectively) than a 1G model ($\chi^2$=1.24). In summary, there is a robust pattern unique to mismatched pC in which it is modeled well with a single continuum of states over all temperatures at which its duplex is intact.

**Supplementary Note 2: Interpretation of $g_F$ spectra obtained through FDCD**

To interpret the FDCD spectra, we assume for simplicity that each conformation *i* is described by a single energy transfer (ET) donor "n" (for "native base") with ET efficiency $a_i$, and that all ET is intramolecular ($c_i = c_j$). This collapses the impacts of individual native bases into collective values of $\Delta\varepsilon_n$ and $\varepsilon_n$ that contain contributions from all bases that transfer energy to pC. The quantities $\Delta\varepsilon_i$ and $\varepsilon_i$ arise from direct excitation of pC itself when the RNA is in conformation *i* which has fluorescence quantum yield $\Phi_i$. Expression 8 (see methods section) then simplifies to:

$$\sum_i \phi_i \left( c_i \epsilon_i + \sum_j a_{ij} \epsilon_j c_j \right) \rightarrow \sum_i \phi_i c_i \left( \epsilon_i + a_i \epsilon_{n,i} \right)$$

We further assume that the ensemble of structures *i* can be approximated with two subpopulations, stacked ("s") and unstacked ("u"), which is supported by our TCSPC measurements. The fluorescence-detected dissymmetry factor is then:

$$g_F = \frac{\phi_u c_u (\Delta\epsilon_u + a_u \Delta\epsilon_{n,u}) + \phi_s c_s (\Delta\epsilon_s + a_s \Delta\epsilon_{n,s})}{\phi_u c_u (\epsilon_u + a_u \epsilon_{n,u}) + \phi_s c_s (\epsilon_s + a_s \epsilon_{n,s})}$$

And the corresponding quantity from standard CD is:

$$g = \frac{c_u (\Delta\epsilon_u + \Delta\epsilon_{n,u}) + c_s (\Delta\epsilon_s + \Delta\epsilon_{n,s})}{c_u (\epsilon_u + \epsilon_{n,u}) + c_s (\epsilon_s + \epsilon_{n,s})}$$

It is helpful to consider these quantities separately in the long-wavelength region of the spectrum (>300 nm) where only pC absorbs, and in the 200-300 nm region where absorbance by the native bases (if they are present) dominates. Beyond 300 nm, the extinction coefficients of the native bases are negligible, yielding:

$$g_F = \frac{\phi_u c_u \Delta\epsilon_u + \phi_s c_s \Delta\epsilon_s}{\phi_u c_u \epsilon_u + \phi_s c_s \epsilon_s}$$

and

$$g = \frac{c_u \Delta\epsilon_u + c_s \Delta\epsilon_s}{c_u \epsilon_u + c_s \epsilon_s}$$

Comparing these two equations, it is evident that $g_F$ can be larger than g in the long-wavelength region if the species (u or s) with the larger quantum yield has the larger CD signal ($\Delta\varepsilon_{u/s}$) and/or the smaller extinction coefficient ($\varepsilon_{u/s}$). $\Delta\varepsilon_s$ will generally be larger than $\Delta\varepsilon_u$ due to close positioning of the bases in a chiral geometry, so a larger $g_F$ than g suggests that stacked conformations have higher fluorescence quantum yield.

Now considering only the contribution of native base absorption followed by ET (dominant at shorter wavelength), we obtain:

$$g_F = \frac{\phi_u c_u a_u \Delta\epsilon_{n,u} + \phi_s c_s a_s \Delta\epsilon_{n,s}}{\phi_u c_u a_u \epsilon_{n,u} + \phi_s c_s a_s \epsilon_{n,s}}$$

and

$$g = \frac{c_u \Delta\epsilon_{n,u} + c_s \Delta\epsilon_{n,s}}{c_u \epsilon_{n,u} + c_s \epsilon_{n,s}}$$

Comparing these equations, it is evident that $g_F$ can be larger than g in the short-wavelength region if the species with the higher product of quantum yield and ET efficiency $\Phi_{u/s}a_{u/s}$ has the larger native base CD signal ($\Delta\varepsilon_{n,u/s}$). Thus, a larger $g_F$ than g at short wavelength suggests that stacked structures have a larger value of $\Phi_{u/s}a_{u/s}$. This product represents the probability of pC emission conditional on absorption by the native base(s). It is reasonable to assume that $a_s$ will generally be larger than $a_u$ due to closer positioning of the bases, so $g_F$ could be larger than g at short wavelength even if $\Phi_u \geq \Phi_s$. The trinucleotide UpCU provides an example of the situation $\Phi_u \approx \Phi_s$ (based on g and $g_F$ being roughly equal >300 nm) and $\Phi_u a_u < \Phi_s a_s$, (based on $g_F$ being larger than g <300 nm), which implies that $a_u < a_s$, as expected. Likewise, the long-wavelength FDCD spectrum of pCG suggests that $\Phi_u > \Phi_s$ while the short-wavelength FDCD indicates $\Phi_u a_u \approx \Phi_s a_s$, together implying $a_u < a_s$.
